# BCR-induced protein dynamics and the emerging role of SUMOylation revealed by proximity proteomics

**DOI:** 10.1101/2020.09.29.318766

**Authors:** Luqman O. Awoniyi, Alexey V. Sarapulov, Sara Hernández-Pérez, Marika Runsala, Diogo M. Cunha, Blanca Tejeda-González, Vid Šustar, M. Özge Balci, Petar Petrov, Pieta K. Mattila

## Abstract

Successful B cell activation, critical for high-affinity antibody production, is controlled by the B cell antigen receptor (BCR). However, we still lack a comprehensive protein-level view of the very dynamic multi-branched cellular events triggered by antigen binding. Here, we employed APEX2 proximity biotinylation to study antigen-induced changes, 5-15 min after receptor activation, at the vicinity of the plasma membrane lipid rafts, wherein BCR enriches upon activation. The data reveals dynamics of signaling proteins, as well as various players linked to the subsequent processes, such as actin cytoskeleton remodelling and endocytosis. Interestingly, our differential expression analysis identified dynamic responses in various proteins previously not linked to early B cell activation. We demonstrate active SUMOylation at the sites of BCR activation in various conditions and report its functional role in BCR signaling through Akt and MAPK axes.

## Introduction

B lymphocytes constitute a critical branch of the adaptive immune system by differentiating into antibody-producing plasma cells after the specific recognition of antigens via their distinctive B cell receptor (BCR). BCR signaling is a robust trigger that leads to phosphorylation of the downstream kinases and cellular structural processes, like actin cytoskeleton reorganisation and internalisation of the BCR, within minutes of activation. Numerous studies on BCR signaling have sketched a picture of a multibranched signaling network that not only triggers cascades to change transcriptional programming but also to alter other cellular machineries such as cytoskeleton reorganisation, endocytosis and vesicle transport, as well as protein degradation (*Harwood & Batista*, 2010; *Kuokkanen et al*., 2015; *Kwak et al*., 2019). Coordinated engagement of such a wide variety of, often interlinked, cellular pathways have challenged our understanding of the early events of B cell activation. One of the major limitations has been the inability to broadly, yet with sufficient spatial and temporal resolution, study the combinations of cellular changes. An efficient way to analyse the signaling network and identify novel players in given cellular pathways is proteomics. Quantitative mass spectrometry (MS)-based proteomics has previously been employed to study BCR signaling. For instance, Satpathy and colleagues used affinity purification to study the dynamics of BCR interactions upon stimulation with biotinylated anti-IgG F(ab’)_2_ at different time points (*Satpathy et al*., 2015). However, the sample preparation for traditional proteomic approaches, relying on co-immunoprecipitation or organelle purification, occurs in the test tube and, thus, poses significant challenges in capturing weak or transient protein interactions (*Qin et al*., 2021). Therefore, in recent years, proximity labeling techniques have become a powerful tool for mapping protein-protein interactions in the native cellular environment. These techniques include antibody-based approaches, such as EMARS and SPPLAT, the promiscuous biotin ligases BioID/BirA*, the engineered ascorbate peroxidases APEX, and the variants thereof (*Bosch et al*., 2021; *Qin et al*., 2021; *Samavarchi-Tehrani et al*., 2020). Proximity-based techniques are typically based on in cellulo biotinylation triggered by enzymes, tagged to a compartment or protein of interest, that generate short-lived biotin radicals that mark their immediate molecular environment. Of these techniques, only SPPLAT (selective proteomic proximity labeling using tyramide) has been successfully applied to the study of BCR interactions, using anti-IgM-HRP antibodies to capture the proteins proximal to the BCR clusters in the chicken B cell line DT40 (*Li et al*., 2014). However, because SPPLAT relies on subjecting living cells to HRP-conjugated antibodies, the biotinylation is thus restricted mainly to the extracellular side of the plasma membrane, failing to capture the whole complexity of processes triggered inside the cell. BioID and APEX2, on the other hand, have a biotinylation radius of 10-20 nm, and they have been successfully used to identify both specific protein interactomes, or immediate protein environments, in various intracellular compartments, such as the mitochondrial matrix, mitochondrial intermembrane space and primary cilia (*Bareja et al*., 2018; *Bosch et al*., 2021; *Hung et al*., 2014; *Mick et al*., 2015; *Rhee et al*., 2013; *Roux et al*., 2012). The second-generation version of APEX, APEX2, that achieves efficient biotinylation in 1 minute, is the fastest and most efficient labeling enzyme to date (*Hung et al*., 2016; *Lam et al*., 2014). As a comparison, TurboID, the fastest member of the BioID family, requires 5-10 min (*Chen & Perrimon*, 2017; *Doerr*, 2018). Therefore, the fast labeling kinetics enabled by APEX2 makes it truly powerful in capturing signaling events with high dynamics, such as receptor signaling. For example, APEX2 was recently used in a tour de force of tracking GPCR signaling and internalisation with a high spatial resolution (*Paek et al*., 2017). However, as for any other fusion protein, expression of APEX2 as a fusion partner of a particular signaling protein can prove technically challenging and also potentially compromise the protein function. In such a scenario, targeting the enzyme to the cellular compartment of interest could provide a better readout. Upon antigen binding, BCR is known to shift from fluid, detergent-soluble plasma membrane domains to less fluid, detergent-resistant cholesterol/sphingolipid-rich membrane domains, called lipid rafts, for signaling and internalisation (*Aman & Ravichandran*, 2000; *Cheng et al*., 1999; *Gupta et al*., 2006; *Sohn et al*., 2006; *Hae et al*., 2008; *Stone et al*., 2017; *Varshney et al*., 2016). This transition provides a spatial window for utilising proximity biotinylation to study BCR signaling. In the past, B cell lipid rafts have been isolated for proteomic analysis via cell fractionation methods (*Gupta et al*., 2006; *Mielenz et al*., 2005; *Saeki et al*., 2003). However, the challenging nature of the biochemical fractionation is illustrated by the limited protein identification, with reported protein numbers in these studies varying between 18 and 39. Here, we pioneer the use of APEX2 for tracking the cellular events occurring at the vicinity of the B cell plasma membrane after activation of the IgM BCR with high spatial and temporal resolution. We utilised the well-defined shift of the BCR to the lipid rafts in order to capture the signaling events and immediate cellular responses at 5, 10 or 15 min after IgM cross-linking, with the 1-min resolution window as allowed by APEX2. Our data, containing 1677 high-confidence hits, provides an encyclopaedia of the proteins locating at the vicinity of the B cell plasma membrane and the lipid rafts while at the same time revealing the dynamics therein induced by IgM stimulation. We identify a wealth of previously uncharacterised proteins responding to the IgM signaling. As validation of our data, we verify the translocation of Golga3 towards activated and internalised BCR, as well as show active SUMOylation at the sites of BCR signaling. Using a pharmacological inhibitor, we demonstrate the functional role of SUMO in BCR signaling to AKT and MAPK cascades.

## Results

### Validation of a B cell line expressing lipid raft-targeted APEX2

In order to gain novel, spatiotemporal information about the immediate cellular responses to BCR activation, we decided to employ a proximity labeling enzyme, APEX2, capable of promiscuously biotinylating proteins within a 20 nm range with a 1-min temporal resolution. As BCR has been reported to translocate to the lipid raft regions of the plasma membrane upon activation (*Aman & Ravichandran*, 2000; *Cheng et al*., 1999; *Gupta et al*., 2006; *Hae et al*., 2008; *Sohn et al*., 2006; *Stone et al*., 2017; *Varshney et al*., 2016), we decided to fuse APEX2 with a specific 7 amino acid lipidation sequence, MGCVCSS, to target it to the lipid raft domains (Fig. 1A). The MGCVCSS sequence contains one myristoylation (Gly-2) and two S-acylation sites (Cys-3 and Cys-5) for palmitoylation, originally identified in the NH2-terminus of Lck, to the NH2-terminus of APEX2. These modifications are responsible for the localisation of Lck to the lipid rafts (*Yasuda et al*., 2000) and target fusion proteins to the membrane, and to the immunological synapse of T cells (*Bécart et al*., 2008; *Bi et al*., 2001). In addition, we equipped our APEX2 construct with an mCherry fluorescent protein to facilitate the detection of APEX2 expression (Fig. 1A, zoom inset).

**Figure 1:**
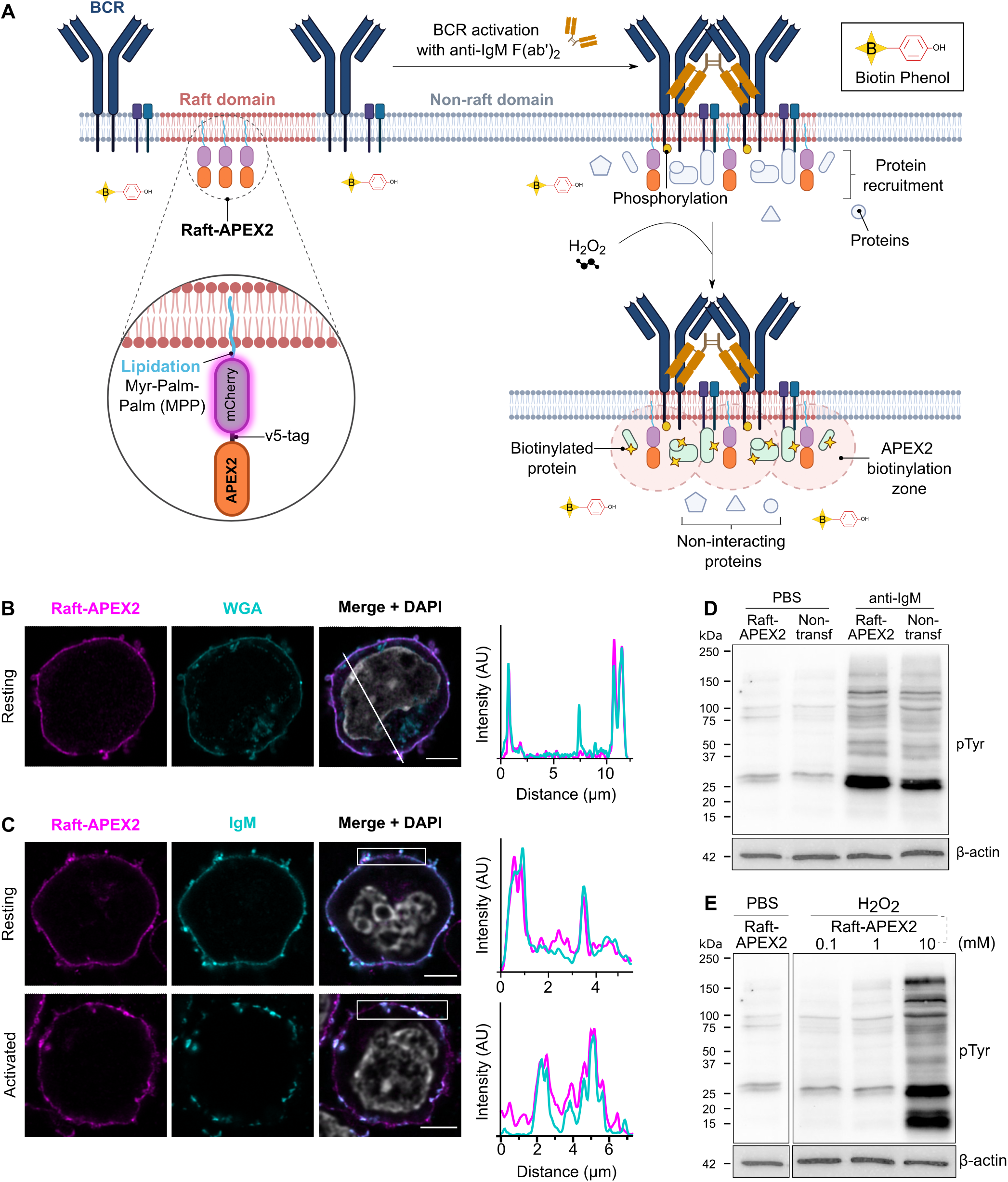
Targeting of APEX2 to the lipid rafts to study B cell activation: design and validation. **A)** Schematic illustration of the raft-APEX2-mediated proximity biotinylation as a read out of BCR signaling response. MPP-mCherry-APEX2 (raft-APEX2) construct is targeted to the lipid raft membrane domains, where also BCR translocates upon activation, and where it induces biotinylation of proteins in the < 20 nm range in a 1 min time window. The technique allows identification and label-free quantification of proteins proximal to the lipid rafts at different time points of BCR activation. **B)** A20 D1.3 B cells expressing raft-APEX2 were settled on a fibronectin-coated glass coverslips for 1 h prior to fixation. Cells were stained with WGA as a membrane marker and DAPI for nucleus. Left: Airyscan super-resolution confocal microscopy was used to image mCherry to visualise raft-APEX2 (magenta), WGA-Atto488 (cyan) and DAPI (grey, merge). Right: line scan analysis of the colocalisation of raft-APEX2 and WGA. **C)** A20 D1.3 B cells expressing raft-APEX2 were settled on a fibronectin-coated glass coverslips for 1 h, activated (bottom panel) or not (upper panel; resting) by addition of 1 μg of HEL for 5 min, and fixed. The samples were stained for IgM and DAPI was used to label the nucleus. Airyscan super-resolution confocal microscopy was used to image mCherry to visualise raft-APEX2 (magenta), IgM (cyan) and DAPI (grey, merge). Right: box scan analysis of the colocalisation of raft-APEX2 and IgM. **D)** Parental (non-transfected) and raft-APEX2-expressing A20 D1.3 B cells were stimulated with 10 μg of anti-IgM F(ab’)_2_ for 10 min and subjected to Western blotting. The membranes were probed with HRP-anti phospho-Tyrosine antibodies, and anti /*beta*-actin as a loading control. **E)** Raft-APEX2 expressing A20 D1.3 B cells were treated with 0, 0.1, 1 and 10 mM H_2_O_2_ for 1 min. Cells were lysed and subjected to Western blotting as in D. See Suppl. Fig. S1C for the uncropped image corresponding to D and E.

Next, we transfected raft-APEX2 into cultured A20 B cells, that stably expressed transgenic hen egg lysozyme (HEL)-specific D1.3 IgM BCR (A20 D1.3) (*Aluvihare et al*., 1997), and generated a stable A20 D1.3 raft-APEX2 cell line. Flow cytometry analysis confirmed that mCherry+/IgM+ cells composed > 99% of the resulting cell line (data not shown). To verify that raft-APEX2 indeed targets to the plasma membrane, we analysed its localisation using AiryScan confocal microscopy (*Huff*, 2015) to gain sufficient resolution to unambiguously detect signals deriving from the B cell plasma membrane. Raft-APEX2 clearly colocalised with the membrane marker, wheat germ agglutinin (WGA), demonstrating strong enrichment at the plasma membrane (Fig. 1B). The lipid raft domains in resting cells are typically very small and transient in nature, making their detection highly challenging even with modern microscopy techniques (*Gupta & DeFranco*, 2003; *Sezgin et al*., 2017; *Stone et al*., 2017). Upon activation, BCR forms clusters that are rich in signaling activity and, at the same time, represent larger detergent-resistant membrane domains (*Gupta & DeFranco*, 2003; *Mattila et al*., 2016; *Stone et al*., 2017). Thus, we next activated the IgM BCR by HEL antigen and followed the colocalisation of IgM with raft-APEX2. As expected, upon cell activation, we detected increased clustering of the IgM as well as enrichment of APEX2 in the same structures (Fig. 1C), indicative for localisation of the raft-APEX2 probe in the IgM signaling clusters. However, a fraction of raft-APEX2 was also visible outside of the IgM clusters.

To investigate the lipid raft domain localisation of raft-APEX2 in more detail, we adopted a flow cytometry-based assay (*Gombos et al*., 2004). We expressed in A20 D1.3 cells raft-APEX2 and model proteins resident either at detergent-resistant (*Martinez-Outschoorn et al*., 2015) or detergent-soluble membrane domains (TDM-GFP; (*Nikolaus et al*., 2010)). In this assay, the cells were treated with 0.1% Triton X-100 to release the detergent-soluble proteins from the plasma membrane, and a detergent resistance index was calculated based on the fluorescence before and after detergent treatment (Suppl. Fig. S1A). The analysis showed that raft-APEX2 resisted the detergent treatment to a similar level with Caveolin-1 (Suppl. Fig. S1B). The detergent-soluble model protein TDM-GFP, on the other hand, was almost completely removed from the plasma membrane after detergent incubation. This analysis provided important support to our approach to use raft-APEX2 as a proxy to label proteins enriched at the lipid raft membrane domains and the vicinity of the signaling IgM.

We then proceeded to test for possible adverse effects caused by the expression of raft-APEX2 in our system. We found normal levels of IgM BCR in our raft-APEX2 cell line, and the receptor was internalised at normal kinetics upon receptor stimulation (data not shown). Furthermore, upon BCR-activation the cells showed indistinguishable levels of phosphorylation, detected by anti-phospho-Tyrosine antibodies, as compared to the parental cell line (Fig. 1D and Suppl. Fig. S1C).

Triggering of the biotinylation activity of APEX2 requires addition of H_2_O_2_. Notably, H_2_O_2_ has been shown to inhibit protein phosphatases and thereby to be able to trigger signaling (*Reth*, 2002; *Wienands et al*., 1996). However, we detected no increase in general protein phosphorylation upon incubation of cells with 1 mM H_2_O_2_ for 1 min, the conditions used to trigger biotinylation by APEX2, while 5-10 times higher concentration of H_2_O_2_ induced profound signaling, consistent with previous reports (*Reth*, 2002) (Fig. 1E and Suppl. Fig. S1C, D). This data suggests that no significant unspecific phosphorylation is triggered by H_2_O_2_ in our experiment settings.

### Proteomic analysis of the lipid raft microenvironment identifies 1677 proteins

For preparing the proximity biotinylation samples, biotin-phenol supplemented cells were activated with potent surrogate antigen, F(ab’)_2_ fragments of anti-IgM antibodies for 0, 5, 10 or 15 min (Fig. 2A). The biotinylation was triggered by adding 1 mM H_2_O_2_ and quenched with the addition of Trolox after 1 min. The biotinylation efficiency was verified in each set of samples by flow cytometric analysis, which typically showed biotinylation in ≈70% of cells (Suppl. Fig. S1E). Lysed cells were subjected to streptavidin affinity purification to pull down the biotinylated proteins for mass spectrometry (MS) analysis (Fig. 2B). To assess for the baseline activity of APEX2 and the contribution of endogenous biotinylation, control samples without H_2_O_2_ (Ctrl 1) or without biotin-phenol (Ctrl 2) were prepared. All conditions were performed in triplicates. Trypsin-digested samples were analysed by LC-ESI-MS/MS using nanoflow HPLC system (Easy-nLC1200, Thermo Fisher Scientific) coupled to the Orbitrap Fusion Lumos mass spectrometer. Peptide/protein calling was done with MaxQuant software (Suppl. File 1A) (*Cox & Mann*, 2008). Differential analysis was done using NormalyzerDE (*Willforss et al*., 2019). After filtration of known contaminants and background, we found 1677 proteins with 2 or more unique peptides identified (Fig. 2C and Suppl. File 1B). High confidence hits from all experimental conditions together are listed in the Supplementary File 1B. As expected, we detected, with very high intensity values, several proteins associated with lipid rafts and BCR signaling (Suppl. Files 2, 4A and Fig2D, E). At the same time, the large total number and variety of identified proteins illustrate the high efficacy of APEX2-mediated protein biotinylation that also reaches also to the cytosolic environment immediately beneath the membrane.

**Figure 2:**
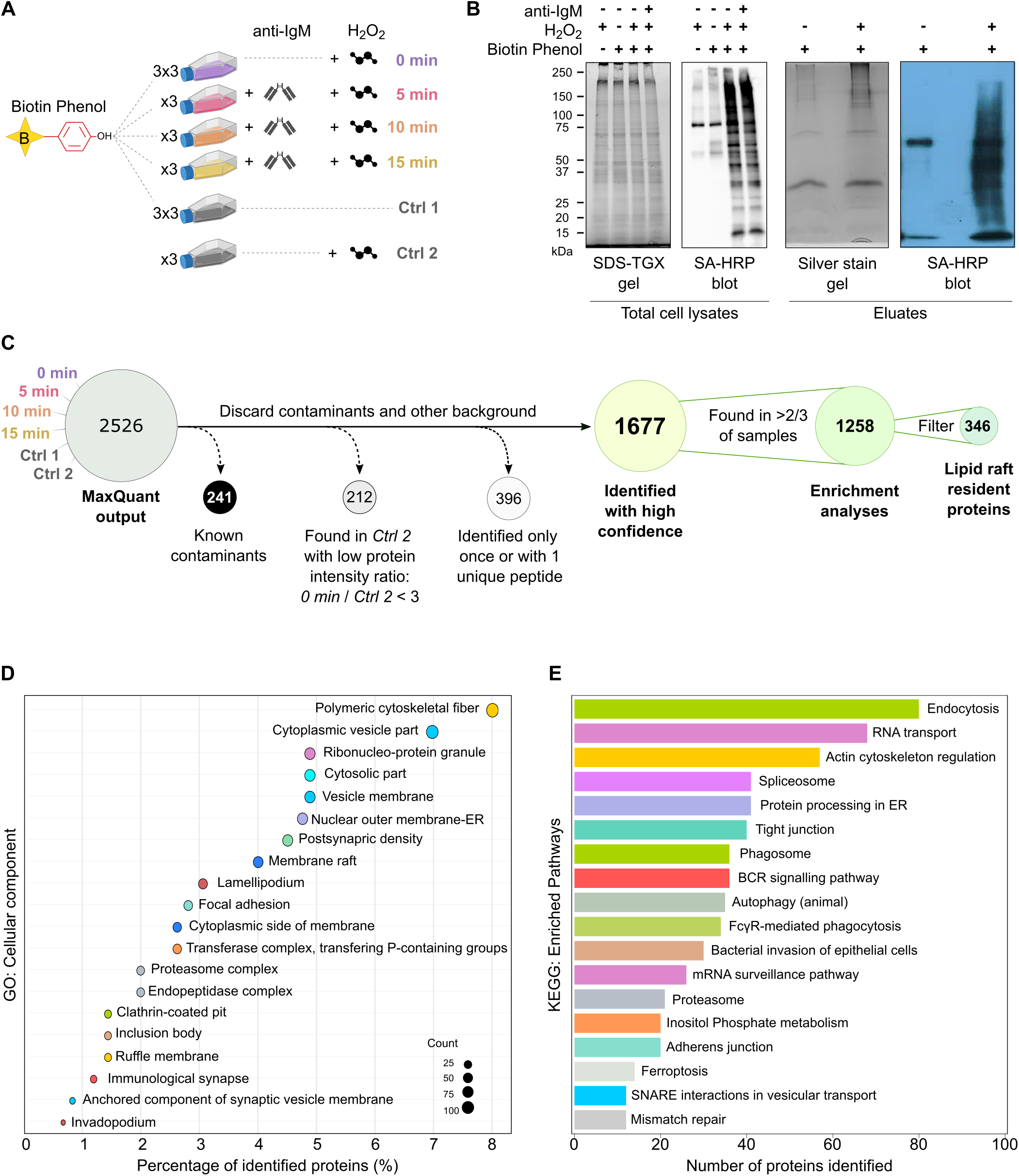
Experimental design and pathway analysis. **A)** schematic representation of the experimental samples and controls prepared and analysed by quantitative label-free MS in this study. Three different biological replicates were prepared for all conditions. **B)** To monitor for basal levels of biotinylation, raft-APEX2 expressing A20 D1.3 B cells were subjected to alternating conditions of anti-IgM stimulation (10 μg/ml, 10 min), H_2_O_2_ (1 mM, 1 min) or biotin-phenol (500 μM, 45 min). Left: Total cells lysates were analysed with TGX Stain-Free SDS gel, and a Western blot probed with streptavidin-HRP. Right: The samples eluted from the streptavidin-coated beads were analysed with a Silver stained SDS-PAGE, and a Western blot probed with streptavidin-HRP. **C)** Flow chart shows the filtering steps used for the data analysis. D) A KEGG pathway enrichment analysis for the 1677 identified proteins shows cellular pathways enriched among the identified proteins. To remove redundancy in the identified pathway terms, terms that had ≥ 50% similarity were grouped and the one with the lowest adjusted p-value is shown. **E)** Classification of the 1677 identified proteins based on cellular component gene ontology (GO) terms. To remove redundancy in the GO terms, GO terms that had ≥ 50% similarity were grouped and the one with lowest adjusted p-value is shown.

### Lipid raft-resident proteins feature stable localisation at the rafts

As a way to further validate the lipid raft localisation of our raft-APEX2 construct, we first short-listed our data for proteins that were likely to reside at the closest vicinity to raft-APEX2. APEX2 exhibits basal activity that leads to low-level release of biotin radicals without added H_2_O_2_, with the aid of low-level endogenous H_2_O_2_. Using a similar approach as Paek and colleagues (*Paek et al*., 2017), we took an assumption that the proteins locating at the closest vicinity to the raft-APEX2 are prone to biotinylation already before extracellular addition of H_2_O_2_. Therefore, the same proteins are expected to get very efficiently labelled upon the addition of H_2_O_2_ and, thus, yield high peptide intensities in the MS data. Thus, we analyszed the protein intensities between the experimental samples and the control samples without added H_2_O_2_ (Ctrl 1). From the proteins identified in Ctrl 1, we further selected those that showed a notable ≥ 1.5 log_2_ fold change upon triggering of biotinylation by H_2_O_2_. Additionally, we filtered out proteins that responded with ≥ 1.0 log_2_ fold change to IgM-stimulation to selectively shortlist only those that were constantly at the closest vicinity of raft-APEX2. As a result, we identified 346 proteins that we considered as B cell raft-resident proteins (Suppl. File 2). Furthermore, almost 90% of the proteins were also found in the available RaftProt databases (*Mohamed et al*., 2019) (Fig. 3A, Suppl. Fig. S2A), providing necessary confidence in the preferred raft localisation of our APEX2 construct. Among the strongest raft-resident proteins that showed most prominent intensity increase upon H_2_O_2_ addition were, for instance, Dock8 and Hcls1, reported regulators of BCR signaling and B cell activation (*Hao et al*., 2004; *Randall et al*., 2010), as well as filamin and spectrin, scaffold proteins linking the plasma membrane and the underlying cytoskeleton (*Liem*, 2016; *Razinia et al*., 2012) (Fig. 3B). An earlier study by Saeki and colleagues identified 34 proteins in the isolated detergent-resistant membrane domains from human Raji B cells (*Saeki et al*., 2003). Out of these 34, our approach identified 20, 10 of which as raft-resident proteins (Suppl. Fig. S2A). The lack of the 10 previously proposed raft proteins could be explained by possible resistance for biotinylation, the differences between the in vitro biochemical fractionation and *in cellulo* labeling, or simply the different cell line used.

**Figure 3:**
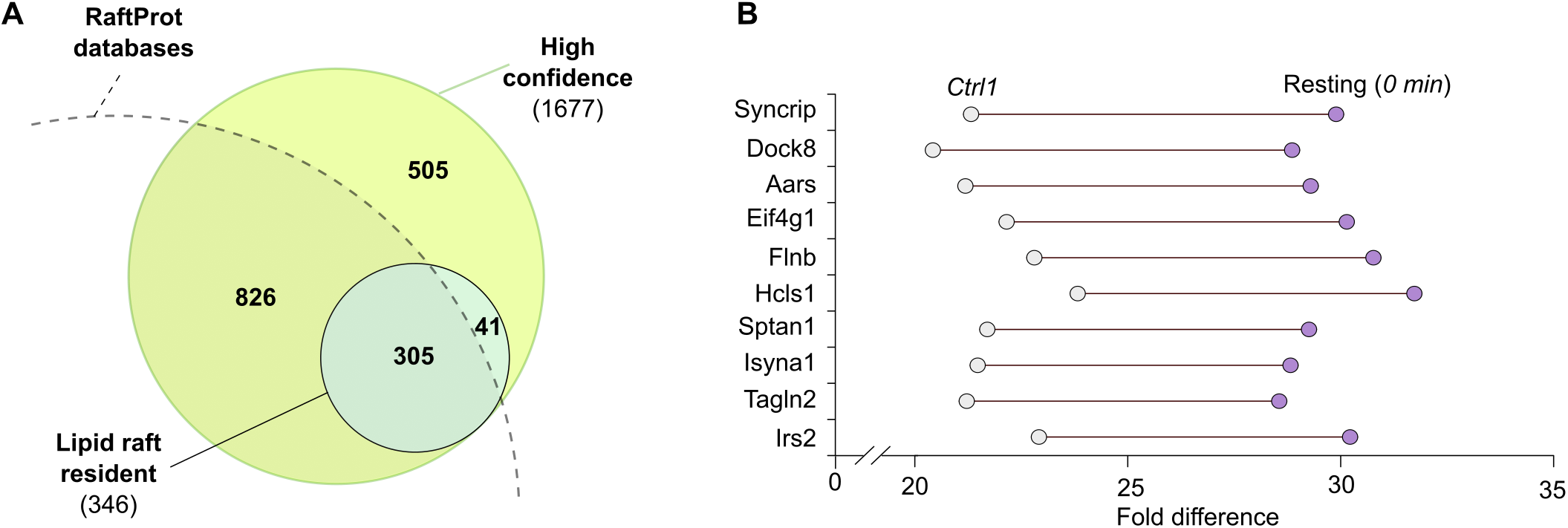
Lipid raft-resident proteins. **A)** A comparison of the lipid raft-resident proteins identified in this study and the whole dataset of 1677 proteins to the RaftProt database shows the overlap between the datasets. **B)** Top ten proteins that show the highest enrichment in the lipid rafts of B cells as reported by raft-APEX2-mediated biotinylation prior (control 1) and post addition (resting, 0 min) of H_2_O_2_. The difference of the protein intensities between the two conditions was used for ranking.

### B cell membrane-proximal proteome reveals a variety of different protein groups

To obtain a broad view on the full set of 1677 proteins identified in our study, we used GO cellular components analysis (*Ashburner et al*., 2000) and KEGG pathway assignment (*Kanehisa et al*., 2016). We identified the highest protein counts in various cytoskeletal and membrane structures, linked to fundamental cell biological pathways such as regulation of the actin cytoskeleton and endocytosis (Fig. 2D, E). Among the more specific terms, we found “Membrane rafts”, “Immunological synapse” and “BCR signaling” to significantly enrich in our data, providing confidence to our approach to detect changes in the protein environment linked to BCR signaling. Out of the total of 1677 high confidence hits in our data (Suppl. File S1B), a large majority, 1143 proteins, were common to all conditions. 48 proteins were specific to resting cells and 40 were specifically detected only upon IgM activation (Fig. 4A, B and Suppl. File 1C, D). Only 2 proteins, Kif20 and Golga3, were identified in all activation time points and not in any of the non-activated controls (Fig. 4A, B and Suppl. File 1D). A significant part of our dataset consisted of proteins that were found present in all or most of the conditions. The protein abundance in each sample was obtained via intensity analysis using MaxQuant software. For statistical analysis of the changes in protein abundance at different conditions, we first applied the criteria of the proteins needing to be present in at least 12 out of 18 experimental samples, which restricted the analysis to 1258 proteins (Fig. 2C,5A, B and Suppl. File 3A). The missing values were then imputed using k-Nearest Neighbor (kNN) and quantitative differential analysis was done using NormalyzerDE (*Willforss et al*., 2019). The majority of the proteins did not undergo significant dynamics upon IgM activation but instead showed relatively stable abundance throughout different conditions. However, 213 proteins showed significant dynamics with log_2_ fold change ≥ 1.5 or ≤ −1.5 upon cell activation (Fig. 5B, Suppl. File S3B). In addition, distinct sets of proteins were found to be enriched or diminished at different time points. While 75, 53 and 55 proteins were significantly altered at 5, 10 and 15 min time points, respectively, only 7 proteins were found significantly altered at all the studied time points (Fig. 5A, B). These findings suggest that while most of the proteins did not dramatically change their localisation, a minor fraction of proteins responded by notable changes regarding their vicinity to the lipid rafts. Our dataset also contained a significant fraction of proteins linked to cellular machineries that are generally not associated with the vicinity of the plasma membrane, such as ribosomes, regulation of translation and RNA transport (Fig. 2E). Arguing against the identification of these proteins as potential background resulting from low-intensity biotinylation or unspecific binding to the streptavidin beads, several such proteins were identified amongst the lipid raft-resident proteins, i.e. with very intimate colocalisation with raft-APEX2 (Suppl. File 2). Interestingly, two earlier independent studies have reported ribosomal and nuclear proteins undergoing S-acylation modification targeting them to the lipid raft domains (*Martin & Cravatt*, 2009; *Yang et al*., 2010). Thus, the prominent lipid raft localisation of ribosomes, could also contribute to the abundance of proteins linked to endoplasmic reticulum, RNA transport and ribonuclear protein granules. Proteins associated with transcriptional regulation, as defined by belonging to the functional category “transcription” at DAVID knowledgebase (*Huang et al*., 2009b,a), constituted about 10% of the detected proteome. We found, for instance, that NF-ϰB proteins, c-Rel, RelA, NF-ϰB2 and regulatory IϰBα are identified in the vicinity of the raft domains of non-activated B cells and show dynamic behaviour upon BCR activation (Suppl. Fig. S3A). FoxO1 transcription factor, whose translocation out of the nucleus is mediated by PI3K-Akt signaling (*Brunet et al*., 1999), associated with raft-APEX2 at 15 min after activation. The transcription factors particularly important for B cell development and differentiation, IRF5 and IRF4, were detected at the raft proximity in all conditions. This finding could be explained by the association of IRFs with membrane-proximal TLR adapter protein MyD88 and Src kinases, including Lyn (*Negishi et al*., 2005; *Ban et al*., 2016). The possible unexpected relationships, suggested by our data, between BCR signaling, plasma membrane, and proteins playing a role in translational regulation, RNA transport and nuclear transport, are attractive topics for future studies.

**Figure 4:**
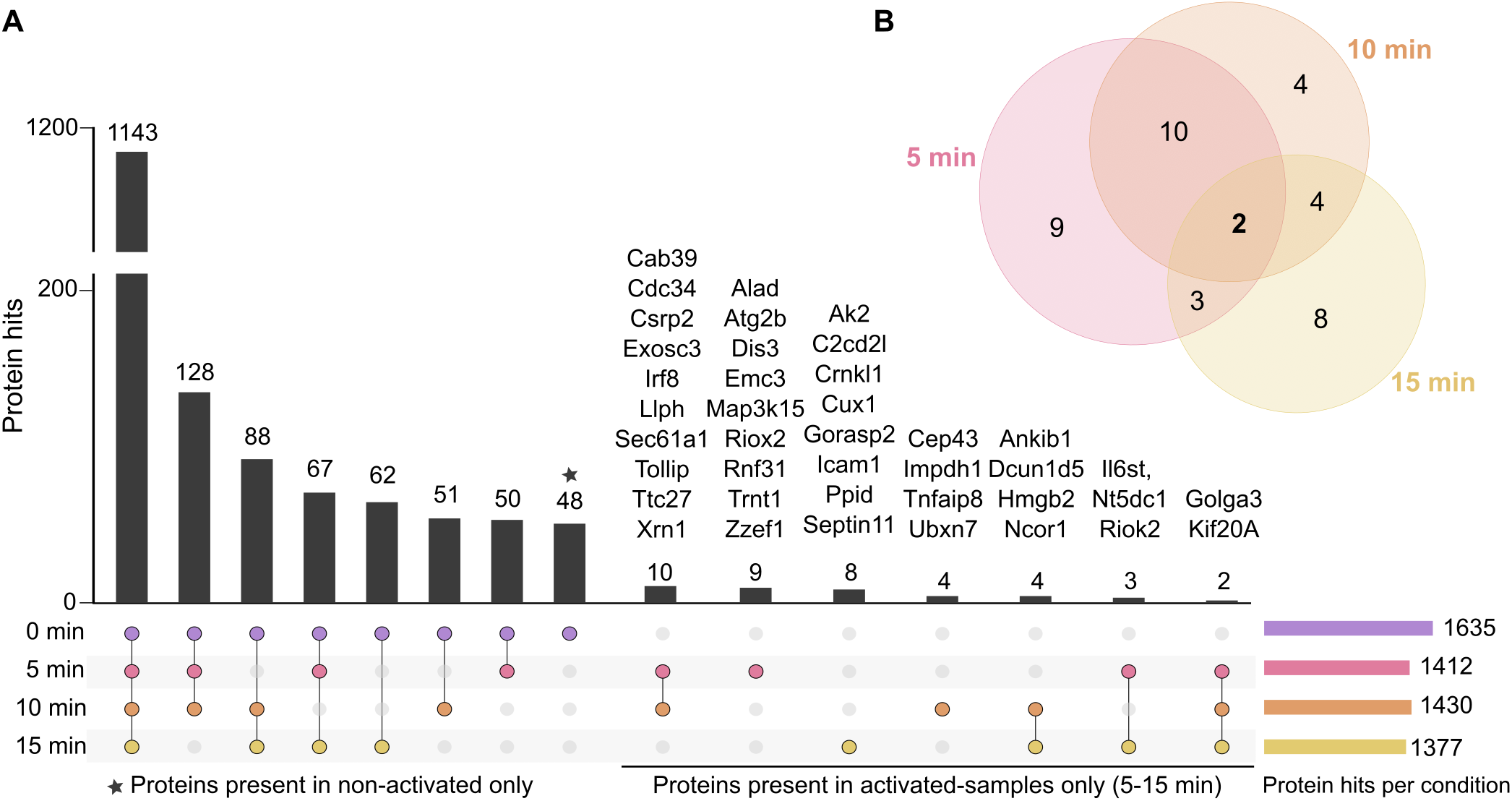
Proteins identified in different conditions of BCR activation. **A)** An UpSet plot showing the numbers of proteins identified in each experimental condition. When 10 or less proteins were identified, the names of the identified proteins are shown on top of the bar. **B)** A Venn diagram showing intersections of proteins identified in activated samples (5, 10 and 15 min of anti-IgM stimulation).

**Figure 5:**
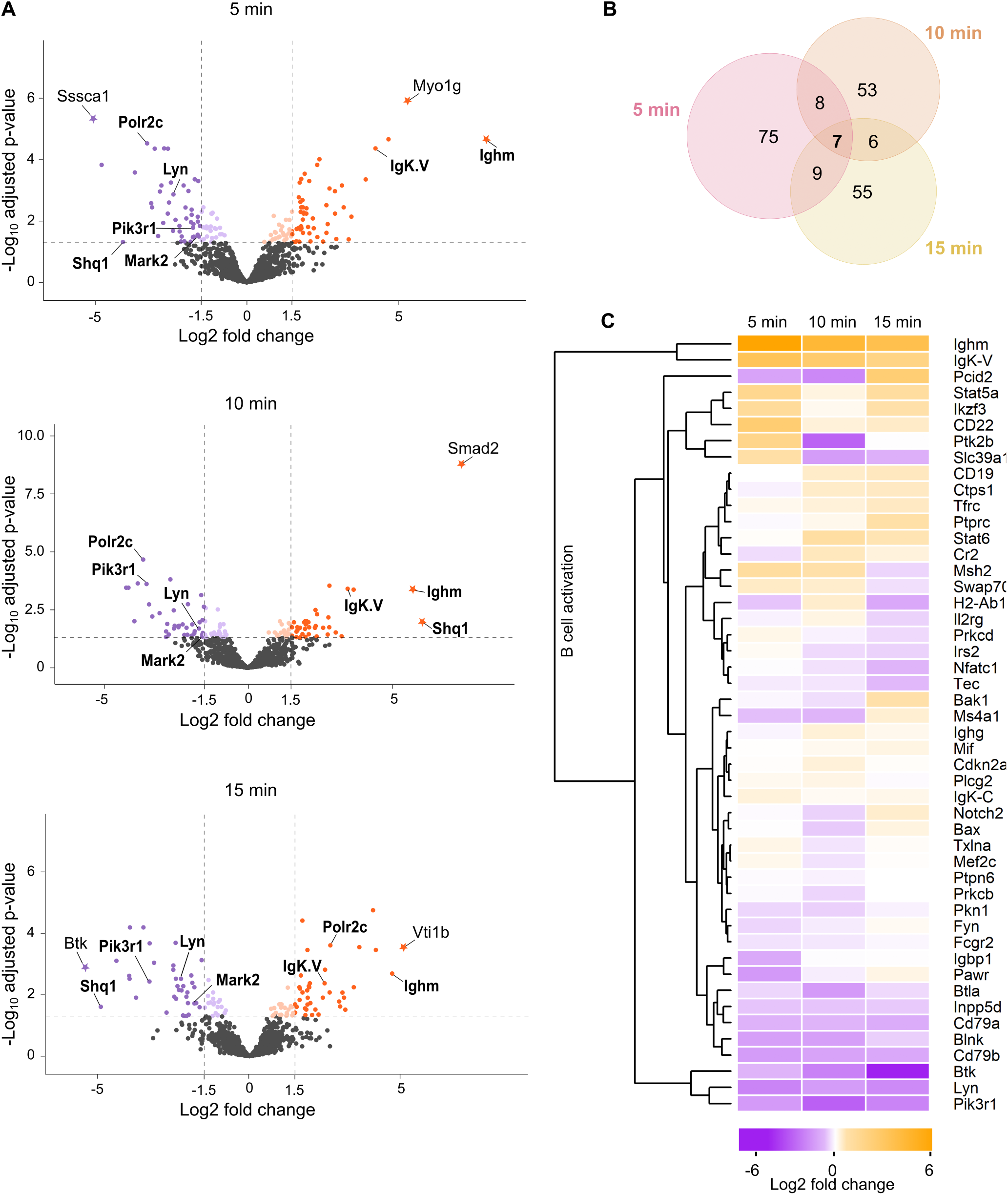
Enrichment analysis. **A)** Volcano plots illustrate the detected protein intensity dynamics upon anti-BCR activation at 5, 10 and 15 min. The data is based on the differential expression analysis of 1258 identified proteins (see Fig. 2C). The proteins showing statistically significant (adjusted p-value ≤ 0.05) enrichment in non-activated conditions are shown in violet and in activating conditions in orange color. The proteins with a log_2_ fold change ≥ 1.5 are further denoted with stronger color tone. The names of the 7 proteins deferentially enriched in all three time points are shown in bold. The proteins with a log_2_ fold change > 5 are denoted with a star symbol. **B)** A Venn diagram showing the numbers of significantly enriched proteins with log_2_ fold change ≥ 1.5 at any time points and their intersections. In total, 213 proteins. **C)** A heatmap of proteins classified to GO term “B cell activation” showing the changes in the protein intensity at different time points of BCR activation.

### The dynamics of proteins linked to BCR signaling

Importantly, Ighm, and IgK-V, the heavy and light chain components of the transgenic IgM BCR specifically stimulated in our setup, showed significantly increased intensity in activated samples (Suppl. File 3A, B; Fig. 5A, C). This finding is in agreement with the well-reported translocation, or induced associated, of IgM to the lipid raft domains. In contrast, Ighg, the endogenous IgG2a BCR, remained essentially unchanged upon receptor activation. This notion proposes that the mechanisms driving activated BCR to the lipid raft domains have specificity to cross-linked receptors and do not carry a notable bystander effect, at least in an inter-isotype manner, that would change the localisation of the unligated BCRs. However, in comparison to IgM, IgG2a was detected at higher levels already in resting cells, suggesting a stronger tendency of this BCR isotype to localise to lipid rafts in resting cells (Suppl. File 2). Interestingly, despite of substantial enrichment of IgM heavy chain and kappa light chain to the lipid rafts upon activation, we saw a decrease in the abundance of Ig*α* and Ig*β*, proteins essential for both signal transmission and the stability and membrane transport of the BCR (Fig. 5C). This finding could be caused by increased shielding of the phosphorylated cytoplasmic tails of the Ig*α* and Ig*β* upon receptor triggering, as a result from the recruitment of the downstream signalosome components. Alternatively, this finding could indicate that the local ratio of Ig*α*/*β* and the BCR heavy chain is not fixed but can be tuned depending on the membrane location and/or the activation state. A tighter molecular organisation could facilitate more efficient sharing of the Ig*α β* sheath molecules upon receptor cross-linking, increasing the IgM:Ig*α*/*β* ratio.

When analysing the known components of the BCR signaling pathway, the results were somewhat unexpected. We did not identify prominent BCR regulator proteins among the proteins exclusively found either in resting cells or activated cells (Fig. 4A), suggesting rather incremental changes in the signaling protein localisation than dramatic translocations induced by IgM engagement. Furthermore, various components of the BCR signaling pathway either did not show significant dynamics, or instead showed diminution while the IgM enriched to the lipid rafts (Fig5C). This finding is consistent with proximal receptor signaling cascades being typically heavily dependent on protein modifications such as phosphorylation, which are not necessarily reflected as changes in total protein localisation.

Among the identified tyrosine kinases involved in early BCR signaling were Lyn, Fyn, Blk, and Btk (Suppl. File 1A). Lyn is considered one of the central kinases triggering BCR signaling, due to its early requirement to phosphorylate Ig*α*/*β* ITAM motifs, and it also preferentially locates to the lipid rafts (*Xu et al*., 2005). Accordingly, we found Lyn in the rafts in all conditions, however, with significantly diminished intensity upon IgM activation. Previous studies, by total internal reflection fluorescence microscopy (TIRFM) of B cells activated on bilayers, indicated that while Lyn does colocalise with clustered BCR, the confinement to the clusters is not very clear. Interestingly, the closest vicinity of Lyn to the BCR, measured by fluorescence resonance energy transfer (FRET), is seen within the first 200 s of BCR activation, after which the interaction diminishes (*Hae et al*., 2008). On the other hand, in the earlier proteomic study, BCR signaling was reported to not affect the enrichment of Lyn in isolated rafts (*Gupta et al*., 2006). The existing ambiguousness over the mechanistic details of Lyn is further complicated by the dual role of the kinase as both negative and positive regulator of BCR signaling (*Xu et al*., 2005). The triggering of the BCR signaling cascade is shared with Fyn, another Src-family protein tyrosine kinase (*Xu et al*., 2005). In our data, Fyn shows higher abundance across the conditions and seems to be largely located at the lipid raftlike regions. Another Src-family kinase identified in our data was Yes. Although not commonly linked to BCR signaling, we detected a significant enrichment of Yes at the lipid rafts at 5 min activation.

From the other components of the BCR signaling pathway, for example, the activatory co-receptor CD19 was constantly found in high abundance in all samples and classified as a lipid raft-resident protein (Suppl. File 2) (Fig. 5C). This data could suggest that the reported enrichment of the CD19 in the BCR signaling microclusters (*Depoil et al*., 2008; *Mattila et al*., 2013) would reflect gathering of the raft domains already containing CD19. Two other transmembrane proteins linked to BCR activation predominantly as negative regulators, Siglec CD22 and FcgR2, were also identified in the data. While CD22 showed enrichment to the lipid rafts at 5 min of activation FcgR2 was defined as lipid raft-resident protein.

Btk, an essential regulator of BCR downstream signaling, also showed increasing strong negative fold-change upon activation, indicating exclusion from the forming BCR clusters in these settings. Btk is known to be recruited to the plasma membrane by PI_(3,4,5)_P_3_ phosphoinositide, a signaling lipid critical for B cell activation (*Saito et al*., 2001). Consistently, we detected a strong diminution of the regulatory subunit of PI3-kinase, Pik3r1, from the lipid rafts upon cell activation. These notions would suggest early separation of the IP_3_-signaling from the immediate vicinity of the BCR. B cell linker (BLNK), a binding partner of Btk and various other BCR signaling proteins showed substantial abundance in the rafts throughout the time points, but also significant downregulation upon BCR activation. Phospholipase-*γ*_2_ (PLC*γ*_2_), which forms the other branch of lipid signaling downstream of BCR, was found constitutively present as a lipid raft-resident protein (Suppl. File 2). In summary, intriguingly, we found BCR signaling proteins mostly either non-dynamically raft-resident, or decreasing from the raft regions upon activation.

### Proximity proteomics identifies multiple vesicle trafficking proteins responding to BCR signaling

In addition to signaling proteins, we detected substantial dynamics of various proteins linked to the steps subsequent to BCR activation, such as cytoskeleton remodelling, endocytosis and membrane trafficking, all essential for internalisation of BCR-antigen complex and further processing for antigen peptide presentation. Our data illuminates the employment of different regulators of these processes, highlighting for example, the existence of various components of the clathrin-mediated endocytosis (Suppl. File 4, Suppl. Fig. S3B). Several of them, such as Cltc, Hip1R and Eps15, were detected as lipid raft resident proteins (Suppl. File 2) or were found differentially enriched during IgM activation (Suppl. File 3), such as AP2 complex subunits alpha and beta. We next sought to validate some of the proteins that were highlighted in our data but not previously linked to BCR signaling. Our attention was drawn to various candidates linked to intracellular membrane traffic that showed specific recruitment towards the lipid rafts upon IgM engagement. For instance, the two proteins identified sorely in activatory conditions, Golgin subfamily A member 3 (Golga3) and kinesin Kif20a (Fig. 4), are both associated with the endomembrane system, but have not previously been associated with BCR signaling. In order to verify the observed dybamics of these proteins, we turned to microscopy. We could not find immunofluorescence-compatible antibodies against mouse or human Kif20a, but went on to visualise Golga3, a poorly understood multifunctional peripheral membrane protein linked to regulation of membrane transport of selected plasma membrane proteins (*Hicks et al*., 2006; *Hicks & Machamer*, 2005; *Williams et al*., 2006), ubiquitination (*Dumin et al*., 2006), apoptosis (*Maag et al*., 2005), and also to dynein function (*Yadav et al*., 2012). In Raji D1.3 human B cells, which were chosen due to better antibodies, we found Golga3 widely distributed in the cells in a vesicular fashion. The distribution of Golga3 vesicles indeed altered upon IgM engagement, and the vesicle pool at the cell membrane became more prominent (Suppl. Fig. S4A). Utilising cell volume segmentation based on microtubule staining (Suppl. Fig. S4C), we selectively analysed the Golga3 vesicles at the vicinity of the plasma membrane before and after BCR activation. We found that the Golga3 vesicles became significantly larger and brighter in the activated cells as compared to the non-activated counterparts (Suppl. Fig. S4D). Also, the shape of the vesicles became more elongated and the vesicles showed notable, yet partial, colocalisation with internalised surrogate antigen. The colocalisation analysis at the cell periphery by Manders’ overlap coefficient showed marked colocalisation of both Golga3 signal in antigen clusters (M1: 0.61), and antigen signal in Golga3 vesicles (M2: 0.56) (Suppl. Fig. S4B. Thus, the immunofluorescence data well supported our proteomics data and showcased Golga3 as a novel protein translocating to the proximity of the BCR upon antigenic activation.

### SUMOylation regulates BCR signaling and immunological synapse formation

To gain a deeper insight on our dataset and uncover more influential proteins or groups of similarly behaving proteins, we performed further bioinformatic analysis on our list of differentially expressed proteins (Suppl. File 3) using unsupervised machine learning and k-means clustering with protein expression log_2_ fold-changes as predictors (*Gu et al*., 2016; *Hartigan & Wong*, 1979). K-means clustering algorithm grouped the proteins into 24 different groups (Suppl. Fig. S5), supporting diverse activation of multiple cellular processes, which we classified based on the major GO biological processes enriched in each group using DAVID (*Jiao et al*., 2012). We got interested in the second largest group, group number 10, containing proteins with gradual increase in their enrichment and classified to be involved in protein transport, nuclear transport, and SUMOylation. Specifically, in this group, RanGAP1 and Small ubiquitin-like modifier 1 (SUMO1) complex with RanBP2 SUMO E3 ligase, also identified in our dataset, critical for nuclear transport of RNAs and proteins (*He et al*., 2021; *Okamura et al*., 2015). Notably, post-translational modification of proteins by addition of SUMO moieties regulates a wide variety of cellular functions, such as DNA repair and replication, signal transduction, cell division, and cell metabolism (**?***Courtois & Fauvarque*, 2018). In our complete APEX2 biotinylated dataset, we identified 9 proteins linked to SUMOylation (Suppl. Fig. S6A). To ascertain a possible involvement of SUMOylation in B cell activation, we went on to investigate the localisation of SUMO upon BCR activation. We first activated A20 D1.3 B cells with fluorescent anti-IgM F(ab’)_2_ for 5, 10, 15 and 30 min and performed immunofluorescence staining for SUMO1. We observed a very strong signal of SUMO1 in the nucleus, reflecting its prominent nuclear functions. Yet, we indeed observed a clear colocalisation of IgM BCR and SUMO in the BCR clusters in the cell periphery, especially in early time points of 5-15 min after activation (Fig. 6A). After 30 min of activation, when the large part of the internalised antigen is clustered to the antigen processing compartments (*Hernández-Pérez et al*., 2020) the colocalisation with SUMO was notably reduced and only some peripheral antigen clusters still showed colocalisation. We were also able to see colocalisation of BCR and SUMO1 in primary B cells isolated from mouse spleen visible by accumulation of SUMO1 at the sites of polarised BCR accumulation, so-called BCR cap, typical for primary B cells (Fig. 6A). The strong nuclear SUMO1 signal significantly challenged the quantitative colocalisation analysis in the cells activated with soluble surrogate antigen. We next activated the A20 D1.3 B cells with antigen-coated beads, mimicking antigen-presenting surface and immunological synapse formation. Importantly, we again saw clear punctate recruitment of SUMO1 to the activatory site, i.e. the surface of the bead, together with BCRs (Fig. 6B). Importantly, such SUMO1 accumulation was not seen with non-activatory control beads (Fig. 6B), although on fibronectin-coated glass SUMO signal was seen to also decorate some of the BCR clusters especially along the filopodia (Suppl. Fig. S6B). Of note, in the 3D stacks of the cells, we frequently noticed another bright SUMO1 cluster close to the nuclear signal. To study this, we polarised the cells by letting them adhere on coated microscope slides and stained them for PCM, a marker for microtubule organizing center (MTOC). We saw a clear enrichment of SUMO1 signal to the MTOC in B cells regardless of their activatory status (Suppl. Fig. S6C, D). We next went on to test possible functional effects of SUMOylation by utilizing a pharmacological inhibitor of SUMOylation, TAK981, which selectively inhibits the SUMO-activating enzyme (SAE) that catalyses the first step in the SUMOylation cascade (*Langston et al*., 2021; *Lightcap et al*., 2021) and was also found in our dataset (Suppl. Fig. S6A). Inhibition of protein SUMOylation by TAK981 at a concentration of 25 μM for 10–30 min efficiently cleared SUMOylation pattern of proteins visible on immunoblot (Fig. 7A). We first probed for the A20 D1.3 cell spreading on the antigen-coated surfaces, mimicking immunological synapse formation. We saw reduced intensity of filamentous actin stuctures in cells treated with TAK981, at 10 or 15 min after activation, indicating lowered efficiency in forming the synapse. Despite a similar trend, we however, did not detect significant reduction in the overall tyrosine phosphorylation in this set-up (Suppl. Fig. S6E, F). We then probed for the activation of phosphorylation cascades downstream of BCR upon SUMOylation inhibition both in mouse primary B cells and in A20 D1.3 cell line using either soluble antigen activation or activation by surface-tethered antigens, mimicking immune synapse formation. In our Western blot analysis, we found a significant reduction in both pAKT and pMAPK1/2 levels in both soluble and surface-bound activating conditions in primary mouse B cells upon inhibition of SUMOylation (Fig. 7B, C). Of note, phosphorylation of Syk was unchanged suggesting that the effect of SUMO occurs at the level of downstream signal regulation. However, no significant functional defects were seen in A20 D1.3 B cell line, fitting with only small defects detected in the spreading response. Finally, we probed for the efficiency of AKT to phosphorylate its target proteins in the conditions where SUMOylation was inhibited, using an antibody that recognises the phosphorylated forms of AKT substrate signature sequences. Although no statistical significance was reached, we saw indications of reduced AKT substrate phosphorylation upon TAK981 treatment in primary B cells, particularly when activated on surface-tethered antigens (p-value = 0.0524) (Fig. 7D, E).

**Figure 6:**
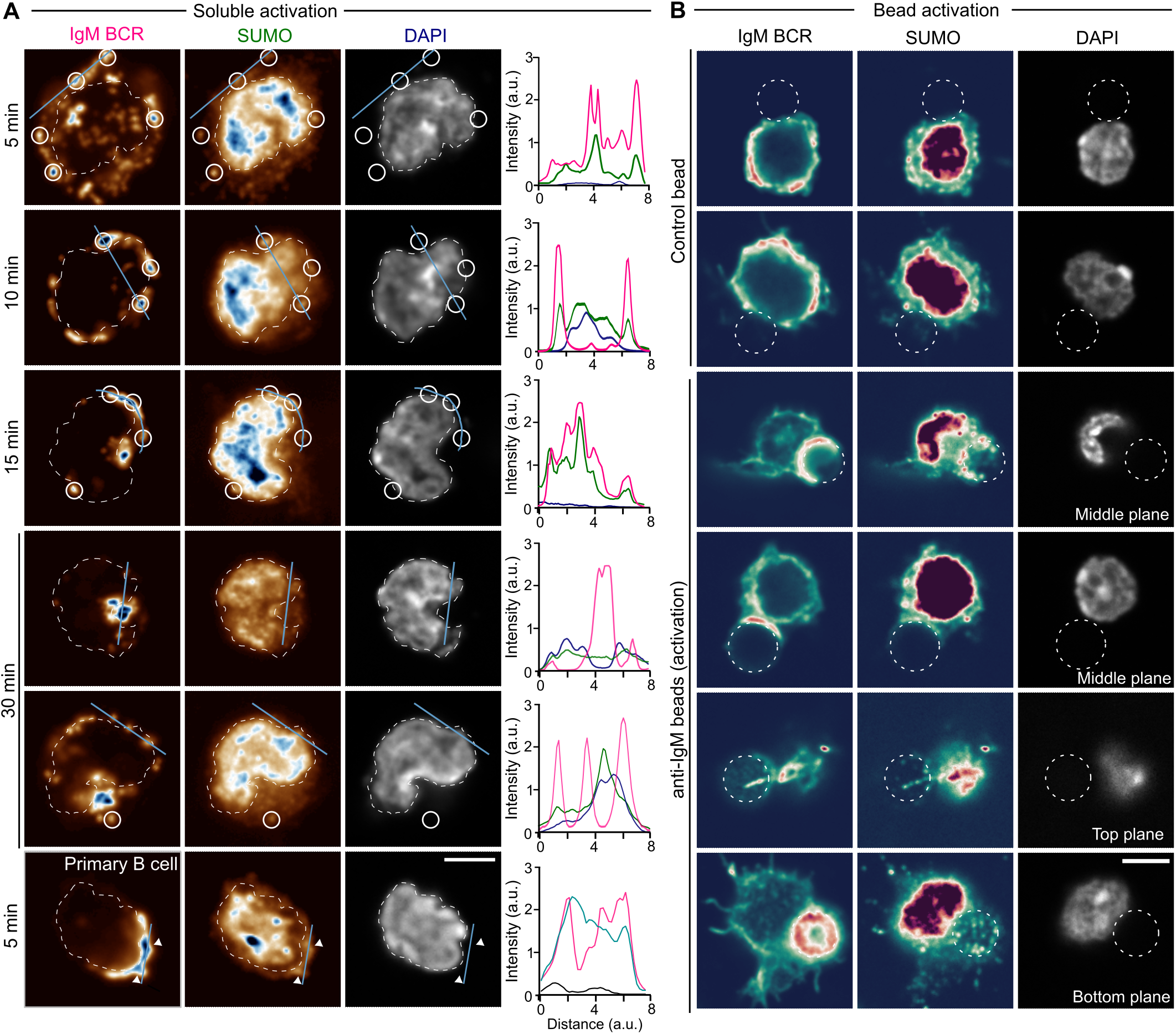
Enrichment of SUMOylation at the sites of BCR activation. **A)** A20 D1.3 cells or primary cells were activated with fluorescently-labelled anti-IgM (soluble surrogate antigen in magenta, 5-30 min), fixed and stained with anti-SUMO (LUT, pseudo-color) and DAPI (grey). The dashed white line outlines the nucleus based on the DAPI staining. The white circles highlight SUMO dots colocalizing with anti-IgM dots. The cyan line represents the line profile shown on the right side of the images (magenta for anti-IgM intensity, green for anti-SUMO1 intensity, and dark blue for DAPI intensity along the line profile). **B)** A20 D1.3 cells were incubated with control beads or anti-IgM coated beads (activators) for 30 minutes, fixed and stained with anti-IgM antibodies (BCR), anti-SUMO and DAPI. The staining are shown using a pseudocolour LUT. The white dashed line shows the position of the bead. Scale bar: 5 *μ*m.

**Figure 7:**
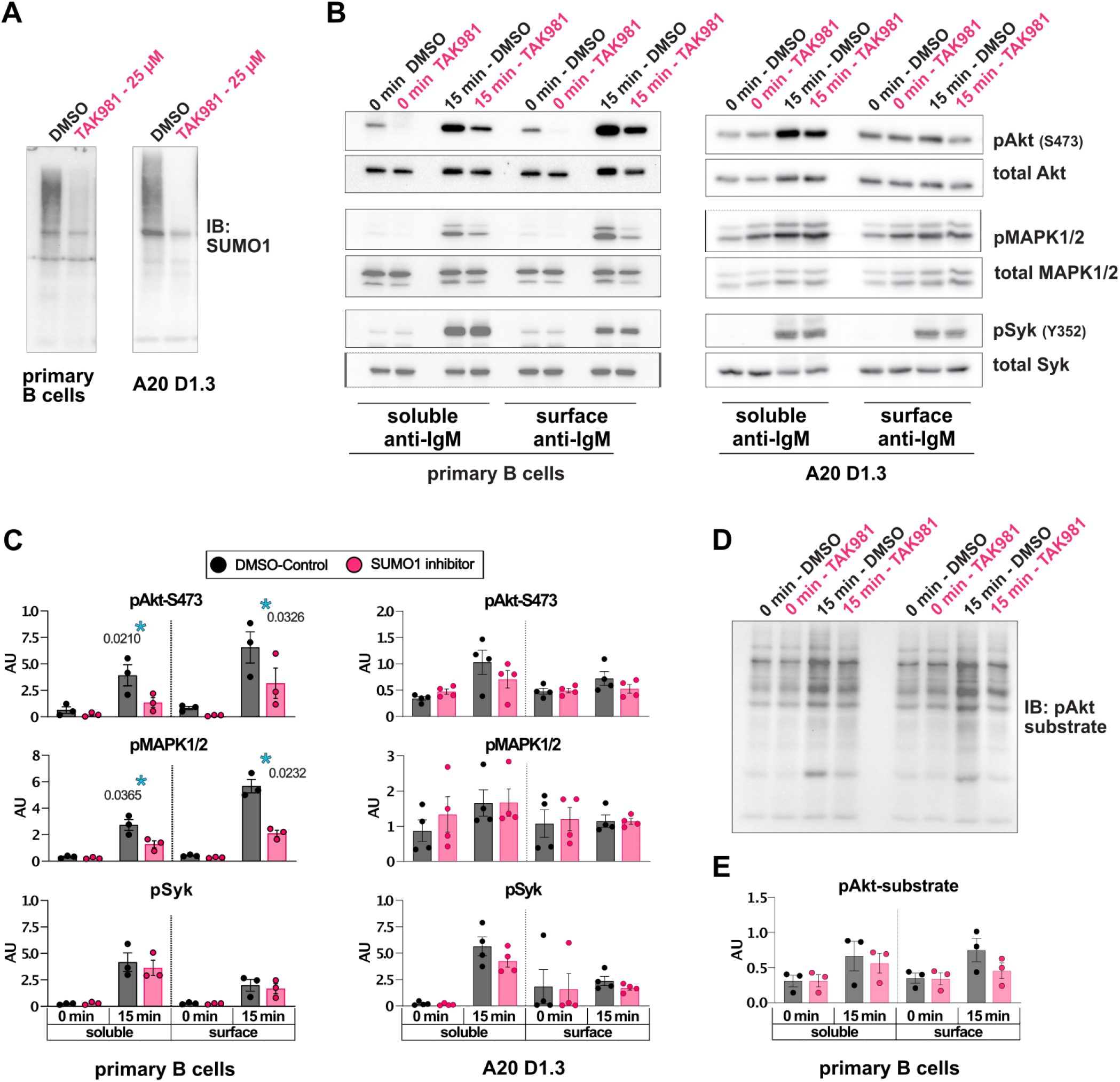
Pharmacological inhibition of SUMOylation leads to reduced AKT and MAPK1/2 signaling downstream of BCR activation. **A)** An immunoblot showing reduction in SUMO1 signal upon 30 min incubation of B cells with 25 μM TAK981. **B)** A20 D1.3 or mouse primary B cells were pre-treated with 25 μM TAK981 for 30 min, stimulated with surface-bound or soluble F(ab’)2 mouse anti-IgM (10 μg/ml) for 15 min in the presence of TAK981 and cell lysates were analyzed for levels of pAkt, pMAPK1/2 and pSyk. **C)** Quantification of the data shown in B. **D)** Primary B cells stimulated as in B were analyzed by immunoblot using pAkt-substrate antibodies recognizing phosphorylated (RXXS*/T*) sequences. E) Quantification of data in D. Data is shown as mean ± SEM from 3-4 independent experiments. *: p < 0.05

## Discussion

To better understand BCR signaling and the immediate, multi-branched cellular responses it triggers, the development of improved large-scale approaches with sufficient spatiotemporal resolution are critical. Here, we pioneer an APEX2-mediated proximity biotinylation proteomics approach, to track at large scale, protein dynamics at the plasma membrane lipid raft domains where BCR translocates upon activation. As APEX2 efficiently biotinylates its vicinity in the range of 20 nm in the time scale of 1 min, it poses significant power to report on protein abundances at a large scale in a time-resolved manner. By identifying and quantitatively analysing over 1600 proteins, we draw a landscape of proteins at or very close to the plasma membrane in B cells and report their dynamics during IgM BCR activation. Furthermore, our data proposes various new protein players responding to the IgM engagement, out of which we validated a vesicle traffic regulator, Golga3, and SUMOylation. We also demonstrate a functional role of SUMOylation on AKT and MAPK activation and show that, in primary mouse B cells, acute lack of SUMOylation during BCR signaling disrupts signal propagation.

With its high efficiency, APEX2-based proximity proteomics provides a sought-after opportunity for an ensemble view of the various cellular machineries triggered upon BCR activation. We faced challenges in fusing APEX2 to the signaling subdomains of the BCR, namely Ig*α* or Ig*β*, and to the proximal signaling protein Lyn, which all failed to get successfully expressed in B cells, while the same constructs were well expressed in fibroblast type of cells. For this reason, we took advantage of the well-described association of the receptor with the lipid rafts induced upon activation. APEX2 targeting by a lipid raft-directed lipidation sequence minimised the risk of interference with the signaling cascades while still reporting about BCR vicinity with reasonable accuracy. Although a fraction of the probe might also remain in the non-raft regions, strong preference to lipid rafts was clear, based on our microscopic and flow cytometric assays (Fig. 1 and Suppl. Fig. S1A, B)). Also, in our shortlisting of the most efficiently and constitutively biotinylated proteins, 90% were found in the RaftProt database (Suppl. File 2). We also saw a drastic enrichment in IgM BCR in activatory conditions (Fig. 5) further proving the correct targeting of the APEX2 and the validity of this approach to report on IgM-proximal proteome.

Of the 1677 proteins identified, a vast majority were detected, at least to some level, both in resting and activatory conditions. This can be a consequence of the known heterogeneity of the raft domains (*Sezgin et al*., 2017) or simply from the high sensitivity of APEX2-mediated biotinylation leading to detection of also the proteins that are present at low levels. A total of 88 proteins were selectively identified either in resting or activated cells (Fig. 4). Additionally, the quantitative analysis revealed 213 proteins with a condition-specific enrichment profile (Fig. 5B). The relatively small proportion of differentially behaving proteins is consistent with only a few changes reported to occur in the isolated lipid rafts upon BCR signaling (*Gupta et al*., 2006). Unfortunately, the study of Gupta and colleagues only reports a selected and limited set of 39 proteins in rafts making thorough comparisons between the data impossible. The highly dynamic nature of the BCR signaling response was clear in the data such that several changes occurring at 5 minutes after activation, for instance, were seen reset by 10 or 15 min (Fig. 4). Interestingly, in general, we found slightly more proteins diminished than enriched at lipid rafts upon signaling. This is in agreement with the fusion and stabilisation of lipid rafts to promote signaling microclusters concomitant with coming together of smaller nanoclusters (*Gupta & DeFranco*, 2003; *Mattila et al*., 2013, 2016; *Stone et al*., 2017), which could reduce the detection of proteins locating preferentially at the borders or surroundings of the rafts. The presence of 48 proteins exclusively in non-activated samples (Fig. 4A and Suppl. File 1B) shows an IgM-induced exclusion of quite a substantial set of proteins, perhaps partially reflecting the reorganisation of the plasma membrane but also suggesting interesting new players orchestrating the signaling cascades. Also, while the abundance of the IgM BCR itself drastically and clearly increased upon activation, the known components of the BCR signaling pathway showed variable responses (Fig. 5C). For example, we noticed an interesting reduction in the abundance of Lyn, Btk, and the Ig*α*/*β* signaling sheath of the BCR, which could indicate that some parts of the signaling pathways separate from the lipid rafts, which on the other hand, become platforms for BCR endocytosis. Such separation of IgM and Ig*α*/*β* has been previously suggested by the finding that Ig*α*/*β* co-precipitate 50-80% less with IgM 15-30 min after receptor cross-linking as compared to the resting conditions (*Vilen et al*., 1999). This finding would also fit with the dissociation-activation model of BCR activation, where opening-up of the BCR oligomers and Ig*α*/*β* sheeth is proposed as a driver of receptor activation (*Yang et al*., 2010). Furthermore, mutagenesis studies on the intracellular tails of Ig*α*/*β* have proposed Ig*α*/*β* signaling and internalisation as mutually exclusive events, such that the phosphorylated Ig*α*/*β* would remain on the cell surface while non-phosphorylated receptors are internalised (*Hou et al*., 2006). However, the reduction of Ig*α*/*β* upon IgM triggering was not reported in the isolated lipid rafts, and no changes in the levels of raft-bound Lyn were shown (*Gupta et al*., 2006). On the other hand, Gupta et al. report a similar reduction in the levels of ezrin and a similar increase in the levels of Myh9, as we observed in our setup. As expected, we found several BCR signaling proteins readily located in the lipid rafts, as suggested by the robust detection of many of them already in the steady-state (Suppl. File 1) as well as the raft-resident, non-BCR-responsive localization of some of them, such as Plc*γ*2 and CD19 (Suppl. File 2). Thus, the translocation of the engaged BCR to the rafts could promote signaling in an energy-efficient manner, as it reduces the need for various other concerted protein translocations.

A crucial role in BCR signaling is played by various protein phosphorylation cascades that have been studied in good detail. Rapidly adjustable post-transcriptional modifications are well-suited drivers for fast signaling events and might not always go fully hand-in-hand with the slower protein translocationschanges in protein localisation, reported by our proteomic study. It is also important to note that while a wealth of our knowledge on BCR signaling comes from studies using soluble antigen, the specific details about signaling protein recruitment are largely derived from microscopy studies using surface-bound antigens, and might differ between different forms of antigenic stimuli (*Depoil et al*., 2008; *Harwood & Batista*, 2010; *Kuokkanen et al*., 2015; *Mattila et al*., 2016). In future studies, it will be very interesting to apply the APEX2 proximity proteomics to the B cell activation by different types of antigens, including surface-bound antigens. Of note, some of the expected proteins linked to BCR signaling, such as Syk, were not identified in the data. Such lack of detection can be caused by several reasons. For example, some proteins are inherently challenging to be detected by MS, or APEX2-mediated biotinylation might be inefficient due to steric obstruction by other proteins, or there might be a lack of suitable amino acid moieties accessible on the protein surface. On the other hand, somewhat unexpectedly, we detected various nuclear proteins, ribosomal proteins and transcriptional proteins. Despite various controls included in our study, some background binding to the streptavidin beads, or biotinylation by the non-localized pool of APEX2 that did not yet reach the cell membrane, is possible. Disputing the possibility of being just unspecific background, many of them showed marked intensity and qualified as B cell lipid raft-resident proteins, or showed significant enrichment upon BCR cross-linking. The detection of many of these proteins could indeed be explained by their direct targeting to the lipid raft membrane domains. Two independent studies, in T cell hybridoma and prostate cancer cells, suggested a set of ribosomal and nuclear proteins to undergo S-acylation and discovered their targeting to the lipid rafts (*Martin & Cravatt*, 2009; *Yang et al*., 2010). Also, increased phosphorylation of eIF3 complex proteins has been observed upon antigen stimulation of B cells (*Matsumoto et al*., 2009), further advocating that some translational regulators could be early targets of BCR signaling. Membrane localisation may serve a regulatory role for these transcriptional and translational regulators and BCR activation with its gathering to the lipid raft domains could, either directly or indirectly, induce the release of this reserve. As we know from previous studies, a large portion of the antigen-BCR complexes are internalised soon after BCR activation, and the complexes are rapidly target to antigen processing compartments (*Hernández-Pérez et al*., 2020). Accordingly, many of the proteins identified with a marked dynamic response to IgM signaling were linked to different branches of intracellular vesicle trafficking or cytoskeletal reorganisation. From the hits not previously linked to BCR signaling, we validated the activation-induced translocation of Golga3 to the vicinity of the plasma membrane and the BCR, as well as a change in the vesicular appearance of Golga3. In the literature, Golga3 has been proposed to recruit cytoplasmic dynein, a minus-end microtubule motor protein, to the Golgi apparatus and to be responsible for the positioning of the Golgi close to the centrosome (*Yadav et al*., 2012). Similarly, Golga3 could be involved in the centripetal movement of internalised antigen vesicles in B cells. As part of a parallel study, we also followed up on another endomembrane trafficking-linked hit from our study, a Q-SNARE Vti1b (vesicle transport through interaction with t-SNAREs homolog 1B), that showed a very strong fold-change in our dataset, being the most enriched protein at 15 min time point (Fig. 5). We characterized its endomembrane localization in B cells and showed its enrichment to the sites of BCR signaling, further supporting the potential of our dataset, although we could not demonstrate a functional role for Vti1b in B cell activation (*Music et al*., 2022). Interestingly, our dataset revealed various proteins linked to SUMOylation pathways. We validated the enrichment of SUMO1 at the sites of BCR activation both upon activation by soluble antigen as well as bead-bound antigen, mimicking immunological synapse formation (Fig. 6). Furthermore, by utilizing an inhibitor for SUMOylation, TAK981, we showed that intact state and dynamics of global protein SUMOylation at the time of BCR triggering contributes to the formation of the B cell immunological synapse and is required for proper phosphorylation of AKT and MAPK1/2, the prominent kinases downstream of BCR activation (Fig. 7). The experiments using an antibody recognizing the phosphorylated consensus sequences of AKT substrates, suggested a reduction in AKT downstream activity upon surface-bound, immune synapse-like, antigen activation. Pronounced downregulation of pAkt and pMAPK1/2 in TAK981-treated normal non-activated WT B cells also suggests that SUMOylation dynamics could be necessary for tonic BCR signaling and may contribute to the previously reported more pronounced depletion of B cells rather than T lymphocytes in mice subjected to multiple exposures of tolerated doses of TAK981 (*Lightcap et al*., 2021). While exhibiting clear enrichment of SUMO-conjugated proteins at the sites of BCR activation, A20 D1.3 B cells, however, seemed functionally less sensitive to SUMOylation inhibition. This indicated a somewhat different wiring of the signaling cascades in the primary B cells versus A20 lymphoma B cells. Altogether, our results draw an important picture of the overall proteome at the B cell plasma membrane and provide both a comprehensive view and unprecedented information on the protein dynamics responding to BCR engagement. In addition to reporting on antigen receptor signaling, our work describes the lipid raft microenvironment in lymphocytes. As lipid rafts have been identified as hotspots for various membrane receptors and signal transduction machineries (*Mollinedo & Gajate*, 2020; *Varshney et al*., 2016), our approach can serve as an easily adaptable platform also for studies of other signaling systems.

## Materials and Methods

### Design and cloning of raft-APEX2

pcDNA3-mito-APEX (*Rhee et al*., 2013), a kind gift from Alice Ting (Addgene plasmid #42607), was used as a template to create and PCR amplify V5 (GKPIPNPLLGLDST) epitope tagged APEX2 cDNA. mCherry with N-terminal seven amino acid sequence (MGCVCSS) that encodes the acylation sequence and APEX2 were then cloned into pcDNA™4/TO plasmid with zeocin selection (Invitrogen V1020-20).

### Cells

The mouse A20 and human Raji B cell lines stably expressing a hen egg lysozyme (HEL)-specific IgM BCR (D1.3) (*Williams et al*., 1994) were a kind gift from Prof Facundo Batista (the Ragon Institute of MGH, MIT and Harvard, USA). A20 D1.3s were maintained in complete RPMI (cRPMI; RPMI 1640 with 2.05 mM L-glutamine supplemented with 10% fetal calf serum (FCS), 4 mM L-glutamine, 50 μM β-mercaptoethanol, 10 mM HEPES and 100 U/ml penicillin/streptomycin). Raji D1.3s were maintained in Raji cRPMI (RPMI 1640 with 2.05 mM L-glutamine supplemented with 10% FCS, 4 mM L-glutamine and 100 U/ml penicillin/streptomycin). Primary B cells were isolated from spleens of 2–3 month old C57Bl/6N mice by negative selection with EasySep™ Mouse B Cell Isolation Kit (# 19854, STEMCELL Technologies).

### Generation of raft-APEX2 expressing stable cell line

Raft-APEX2-pcDNA™4/Zeo/TO plasmid was transfected into A20 D1.3 cells line as previously described (*Šuštar et al*., 2018). In brief, 4 million cells were resuspended in 180 μl of 2S transfection buffer (5 mM KCl, 15 mM MgCl_2_, 15 mM HEPES, 50 mM Sodium Succinate, 180 mM Na_2_HPO_4_/NaH2PO4 pH 7.2) containing 10 μg of plasmid DNA and electroporated using AMAXA electroporation machine (program X-005, Biosystem) in 0.2 cm gap electroporation cuvettes. Cells were then transferred to 4 ml of cRPMI containing extra 10% FCS to recover overnight. Cells were sorted, using Sony SH800 Cell Sorter, single cell/well into 96-well flat bottom plates containing 100 μl of cRPMI containing 20% FCS. Cells were left to recover in cRPMI supplemented with extra 10% FCS for 48 h before adding Zeocin (600 μg/ml final concentration). Clones expressing raft-APEX2 were selected, expanded few weeks after sorting and the expression of raft-APEX2 was verified with flow cytometry analysis for mCherry, V5 and functional biotinylation.

### Proximity biotinylation

10 × 10^6^ A20 D1.3 cells expressing raft-APEX2 were treated with 500 μM biotin-phenol (BP) (Iris-Biotech, CAS no.: 41994-02-9) for 45 min and activated with 0 or 10 μg/ml of goat anti-mouse IgM F(ab’)_2_ fragments (Jackson ImmunoResearch 115-006-020) for 5, 10 or 15 min. 1 mM H_2_O_2_ (Sigma-Aldrich, cat. no. H1009-100ML) was added for 1 min and then quenched with 2X quenching solution (20 mM sodium ascorbate, 10 mM Trolox (Sigma-Aldrich, cat. no. 238813-1G) and 20 mM sodium azide solution in PBS). Cells were repeatedly washed 4 time with 1X quenching solution. Non-biotinylated control samples were prepared similarly but without anti-IgM and H_2_O_2_. Background control samples were prepared similarly but without BP and anti-IgM. To validate biotinylation for each experiment we used flow cytometry, where cells were fixed, permeabilised and stained with streptavidin 633 (Thermo Fischer Scientific). Samples were lysed with modified RIPA buffer (50 mM Tris, 150 mM NaCl, 0.1% SDS, 2% Octyl glucoside (Sigma-Aldrich, 29836-26-8), 0.5% sodium deoxycholate and 1% Triton X-100, pH 7.5) with 1× protease phosphatase inhibitor mini tablet (1 tablet/10 ml, Thermo Fisher Scientific, cat. no. A32961). Lysate concentrations were measured using Pierce 660 nm protein assay (Thermo Fisher Scientific, cat. no. 22660), aliquoted into 360 μg of total protein/aliquot, snap frozen and stored at −80 °C.

### Streptavidin pull-down of biotinylated proteins

350 μg of whole lysate diluted in additional 500 μl of RIPA buffer (50 mM Tris, 150 mM NaCl, 0.1% SDS, 0.5% sodium deoxycholate and 1% Triton X-100, pH 7.5, 1× protease phosphatase inhibitor mini tablet) was incubated with 30 μl (0.3 mg) of streptavidin magnetic beads (Pierce, cat. no. 88817) for 1 h at RT on rotation. Beads were washed several times with 1 ml volumes for 5 min each, on ice, as follows: twice with RIPA buffer, twice with 1 M KCl, twice with 0.1 M Na2CO3, once with 4 M urea in 10 mM Tris-HCl pH 8.0, once with 50 μM biotin, 4 M urea in 10 mM Tris-HCl pH 8.0, and three times with RIPA buffer. Biotinylated proteins were eluted by boiling the beads in 30 μl of 3× SDS loading buffer supplemented with 2 mM biotin and 20 mM DTT for 10 min.

### In-gel Digestion

Eluted samples were run on 10% SDS-PAGE and the gel was stained with SimplyBlue SafeStain (ThermoFisher Scientific, cat. no. LC6065). For the digestion, the protocol adapted from Shevchenko et al. was used (*Shevchenko et al*., 2007). Each gel lane was cut into 4 pieces that were washed twice with 200 μl of 0.04 M NH_4_HCO_3_/50% Acetonitrile (ACN) and dehydrated with 200 μl 100% ACN. Then, gel pieces were rehydrated in 200 μl of 20 mM DTT and dehydrated again as above. Gel pieces were then rehydrated with 100 μl 55 mM Iodoacetamide for 20 min in the dark, RT, washed twice with 100 μl 100 mM NH_4_HCO_3_ and dehydrated as above. 30 μl of 0.02 μg/μl of trypsin (Promega V5111) solution was added to the gel pieces for 20 min followed by addition of 60 μl solution containing 40 mM NH_4_HCO_3_ / 10% ACN to completely cover the gel pieces and the samples were incubated at 37°C for 18 h. Peptides were extracted using 90 μl of ACN followed by 150 μl of 50% ACN / 5% HCOOH at 37 °C for 15 min.

### Mass spectrometry analysis

Data were collected by LC-ESI-MS/MS using a nanoflow HPLC system (Easy-nLC1200, ThermoFisher Scientific) coupled to the Orbitrap Fusion Lumos mass spectrometer (Thermo Fisher Scientific, Bremen, Germany) equipped with a nano-electrospray ionisation source. Peptides were first loaded on a trapping column and subsequently separated inline on a 15 cm C18 column. A linear 20 min gradient from 8 to 39% was used to elute peptides. MS data was acquired using Thermo Xcalibur 3.1 software (Thermo Fisher Scientific). A data dependent acquisition method that consists of comprising an Orbitrap MS survey scan of mass range 300-2000 m/z followed by HCD fragmentation was used.

### Protein Identification

The raw MS data were processed using MaxQuant software version 1.6.0.1 (*Cox & Mann*, 2008). MS/MS spectra were searched against mouse UniProt (reviewed (Swiss-Prot), released September 2019) using Andromeda search engine (*Cox et al*., 2011). The following configuration was used for MaxQuant search: Digestion set to trypsin, maximum number of missed cleavages allowed set to 2, fixed modification set to Carbamidomethyl and variable modifications set to N-terminal acetylation and methionine oxidation. Peptide and protein false discovery rate were set to 0.01. Match between runs was enabled. MaxLFQ that enables determination of relative intensity values of proteins and also normalise proteins intensity between samples was enabled but not used in downstream analysis (*Cox et al*., 2014). After MaxQuant run, 2526 proteins were identified, from which contaminants and reverse hits were removed. For further analysis, only the proteins identified with at least 2 unique peptides were considered identified (1677 proteins). The identified proteins were then classified using both KEGG pathway analysis (*Kanehisa et al*., 2016) and gene ontology (GO) classification (*Ashburner et al*., 2000). The proteomic data set generated in this work will be submitted to PRIDE (*Vizcaíno et al*., 2016).

### Proteomics and differential expression analysis

Normalisation and differential expression analysis were done using NormalyzerDE (*Willforss et al*., 2019) tools in Bioconductor. Quantile normalisation was selected as the best normalisation method following comparison of various normalisation methods in NormalyzerDE (data not shown). Prior to normalisation and differential expression analysis, identified proteins with missing values ≥7 out of 18 conditions were filtered out. For the remaining proteins, missing value imputation was done using k-Nearest Neighbor (kNN) Imputation. Differential expression analysis was done using NormalyzerDE with statistical comparison method set to limma, logTrans set to FALSE, leastRepCount set to 1 and sigThresType set to FDR (Benjamini-Hochberg corrected p-values). To identify proteins that are likely raft-resident a strategy adapted from Paek et al., 2017 was used (*Paek et al*., 2017). We first selected proteins with ≥1.5 log_2_ fold-change in non-activated biotinylated sample compared to control samples (sample without H_2_O_2_ triggered biotinylation). Then, proteins that show log_2_ foldchange ≥ 1 in surrogate antigen stimulated samples compared to non-activated samples were filtered out. The list of the proposed raft-resident proteins was compared with previously published mammalian lipid raft proteins available in RaftProt database (https://raftprot.org/) (*Mohamed et al*., 2019).

### Bioinformatics analysis

All downstream analysis were carried out with R. Enhancedvolcano, which was used to generate volcano plots (*Blighe et al*., 2019). Enrichment analysis was done using R clusterProfiler package (*Yu et al*., 2012). UpSet plot was constructed to show intersect between conditions. A Venn diagram was constructed, using DeepVenn, to depict the intersection of proteins significantly enriched upon BCR cross-linking at different time points. clValid, R package in CRAN was used to determine the number of k clusters prior k-means analysis (*Brock et al*., 2008). The heatmap for different clusters was plotted using ComplexHeatmap R package (*Gu et al*., 2016).

### Western blotting

Raft-APEX2 A20 D1.3 cells were starved for 20 min in serum-free media and incubated with either 10 μg/ml of goat anti-mouse IgM F(ab’)_2_ fragments (Jackson ImmunoResearch 115-006-020) for 10 min or 0.1, 1, 2, 5, and 10 mM H_2_O_2_ for 1 min. After which cells were lysed with 2x SDS PAGE loading buffer, sonicated and proceeded to SDS gel and Western blotting, probed with streptavidin-HRP (Life Technologies, cat. no. S-911). For the analysis of BCR signaling in TAK981-treated B cells, A20 D1.3 or primary splenic B cells were transferred in plain RPMI, incubated with 25 μM TAK981 or DMSO as a control for 30 min, and then stimulated with soluble or pre-adsorbed plate-bound (both at 10 μg/ml) goat anti-mouse IgM F(ab’)2 fragments (Jackson ImmunoResearch 115-006-020) for 15 min in the presence of TAK981 inhibitor and lysed by addition of 5x SDS lysis buffer to the final concentration of 2% SDS. Lysates were sonicated and subjected to SDS PAGE. The blots were processed as described in (*Sarapulov et al*., 2020). Briefly, the blots were probed with antibodies against phosphorylated forms of proteins, stripped and reprobed with antibodies against total corresponding proteins. Following antibodies have been used: anti-pSyk (Y319)/pZap-70 (Y352) (#2701), anti-Syk (#13198), anti-pAkt (S473) (#4058), anti-Akt (#2938), anti-pMAPK1/2 p44/p42 (T202/Y204) (#9101), and anti-MAPK1/2 (#9102), and pAkt-substrate (#9614), all from Cell Signaling Technologies. Inhibition of sumoylation was confirmed with anti-Sumo 1 antibody [Y299] (ab32058, Abcam). Images were background subtracted (ImageJ) and integrated densities for bands corresponding to phospho-specific protein forms were normalized by corresponding total protein densities. Integrated densities of pAkt-substrate signal were quantified from processed images (Enhance contrast… > normalize histograms, 0.001% saturated pixels). Data was analyzed by repeated measures 2-way ANOVA with Šídák multiple comparisons test.

### Detergent Resistance Membrane analysis by flow cytometry

A20D1.3 cells were electroporated with raft-APEX2, caveolin-1-RFP as a raft marker, or GFP fused to influenza virus hemagglutinin transmembrane domain (TDM) (*Nikolaus et al*., 2010) as a non-raft marker. The assay was performed according to Gombos (*Gombos et al*., 2004). In short: 24 hours after transfection, every condition was divided into two tubes. Cells (106/ml in Imaging Buffer) were kept on ice, and Triton X-100 was added to a final concentration of 0.1% to one of the tubes, while the other tube was left untreated, and fluorescence was recorded immediately in a flow cytometer.The parameter of detergent resistance was calculated as DRI=(FL_det_-FLBg_det_)/(FL_max_-FLBg), where FL_det_ stands for fluorescence of the cells treated with detergent for 5 min, FLBg_det_ for autofluorescence of the detergent-treated cells, FL_max_ for fluorescence of labelled untreated cells (proportional to the protein expression level), FLBg for autofluorescence (background) of the unlabelled cells. The experiment was repeated at least 6 times for each marker.

### AiryScan confocal microscopy to analyse raft-APEX2 localisation

Glass-bottom microscopy chambers (Mattek) were coated with 4 μg/ml fibronectin in PBS for 1 h RT and washed with PBS. Similarly to the mass spectrometry samples, cells were incubated in 500 μM BP in complete medium at 37 °C for 45 min. The cells were then let to settle on the microscopy chambers at 37 °C for 15 min and incubated with 1 mM H_2_O_2_ together with 4% paraformaldehyde and 0.1% glutaraldehyde to immediately fix the sample, washed and continued to fix for further 10 min. Cells were washed, blocked in blocking buffer (BSA + goat serum) at RT for 1 h, labelled with Atto488-labelled WGA, washed, permeabilised with 0.1% Triton X-100, at RT for 5 min, blocked again, and labelled with streptavidin-Alexa Fluor® (AF)-633 (1:2000) and DAPI at RT for 1 h. After washing, the samples were mounted in Vectashield. For visualisation of raft-APEX2 upon BCR activation, after settling, 1 μg/ml of HEL antigen was added on ice, then incubated at 37 °C for 5 min and fixed for processing as above with exception of staining with Atto488-labelled goat anti-mouse IgM F(ab’)_2_ fragments (Jackson ImmunoResearch 115-006-020) instead of WGA-Atto488. Images were acquired using a laser scanning confocal microscope LSM880 (Zeiss) with an Airyscan detector (32-channel Airyscan Hexagonal element) equipped with 405 (Diode), 488 (Argon) and 633 (HeNe) laser lines and an oil-immersion 63× Zeiss Plan-Apochromat objective. Images were acquired using the standard super-resolution mode (Zen Black 2.3). The profile intensity analysis was done in Fiji ImageJ (NIH).

### Immunofluorescence sample preparation

Unless otherwise stated, 12-well slides (Thermo Scientific, ER-202-CE24) were coated with 4 μg/ml fibronectin (non-activatory ligand) in PBS for 20 min at RT. A20 D1.3 or primary B cells isolated from C57BL/6 mice spleens were seeded on the fibronectin-coated wells and incubated at 37 °C for at least 15 min to allow adhesion. Then, cells were fixed with 4% PFA for 10 min at RT and blocked/permeabilized for 20 min at RT (5% donkey serum with 0.3% Triton X-100 in PBS). After blocking, samples were stained with primary antibodies for 1h at RT or 4 °C O/N in staining buffer (1% BSA, 0.3% Triton X100 in PBS), followed by washes with PBS and incubation with the fluorescently labelled secondary antibodies for 30 min at RT in PBS. For immunostaining, anti-Sumo1 (Y299, Abcam, ab32058) was used at a dilution 1:500, anti-PCM1 (Santa Cruz Biotechnologies, sc-398365 AF647) at 1:400, Donkey anti-mouse IgM F(ab’)2 - RRX (Jackson ImmunoResearch, #715-296-020) at 1:500 to stain the IgM BCR, and anti-Phosphotyrosine (4G10, Merck Millipore, 05-321) at 1:400. Secondary antibodies anti-rabbit IgG AF488 (Thermo Fisher Scientific A-21206) and anti-mouse IgG2b AF633 (Thermo Fisher Scientific A-21146) were used at 1:500. For actin staining, phalloidin 555 (Acti-stain 555 Cytoskeleton Inc, #PHDH1-A) at 1:400 dilution was used. Samples were mounted using FluoroMount-G containing DAPI (Thermo Fisher, #00495952). Images were acquired on a 3i CSU-W1 Marianas spinning disk confocal microscope (Intelligent Imaging Innovations) equipped with a 63× Zeiss Plan-Apochromat objective (NA 1.4) and a Hamamatsu sCMOS Orca Flash4.0 camera (2048 x 2048 pixels, 1 x 1 binning). For experiments where activation with soluble antigen was needed, A20 D1.3 or primary B cells were labelled on ice for 10 minutes with 10 μg/ml of Alexa Fluor® 647 donkey anti-mouse IgM (#715-605-140, Jackson ImmunoResearch) or RRx goat anti-mouse IgM (#115-295-205, Jackson ImmunoResearch), washed with PBS to remove excess unbound antigen and resuspended in Imaging Buffer (PBS, 10% FCS). Cells were then seeded on the fibronectin-coated slides for 5 to 30 minutes to allow activation and samples were processed as described above. For experiments where activation with surface-bound antigen was used, 12-well slides were coated with 10 μg/ml F(ab’) Fragment Goat Anti-Mouse IgM (Jackson ImmunoResearch, 115-006-020) in PBS for 1 hour at room temperature or fibronectin as a control, as described above. Cells were seeded on the coated slides, activated for 5-15 min, and processed as described above. The profile intensity analysis was done in Fiji ImageJ (NIH) (*Schindelin et al*., 2012). For experiments where cells were activated with antigen-coated beads, 5 μm Streptavidin beads (Bangs Laboratories, # CP01N/10984) were coated with 10 μg/ml of biotinylated goat anti-mouse IgM at 37 °C for 30 min (shaking 1000 rpm) and washed in 2% BSA/PBS. Uncoated beads were used as a negative control. Cells were mixed with the beads (1:1) and plated on the fibronectin-coated wells for 30 min (+37 °C, 5% CO2) to trigger activation. Samples were then processed and imaged as described above. In those experiments were the SUMO1 inhibitor was used, A20 D1.3 cells were suspended in Imaging Buffer and pretreated with the SUMOylation inhibitor TAK981 (MedChemExpress, HY-111789) at 25 μM final concentration or DMSO as a control in the incubator (5% CO2, +37 °C). After that they were seeded in the 12-well slides coated with 10μg/ml F(ab’) Fragment Goat Anti-Mouse IgM (Jackson ImmunoResearch, 115-006-020) in PBS for 1 hour at room temperature and activated for 15 or 5 min in the incubator, being treated with the inhibitor for 45 min in total, and processed as above. Samples were imaged as above and, using Fiji Image J, the cell area was thesholded, based on phalloidin channel, for the analysis of area and signal intensity. Data was analyzed by a paired t-test on the mean values of each experiment.

### Immunofluorescence sample preparation, acquisition and image analysis for Golga3

8-well polymer coverslips (μ-Slide 8, high-well, IBIDI 80806) were coated with CellTak substrate (CellTak, Corning®, purchased from Sigma-Aldrich DLW354242) according to the manufacturer’s recommendations. In short, 80 μl of 56 μg/ml CellTak in H_2_O was applied in each well. CellTak was topped with 120 μl of 0.1 M NaHCO3 pH 8.0. to activate the reaction. The slides were incubated for 1-2 h at RT, washed 1x with H_2_O, dried and stored at +4°C. 150.000 Raji D1.3 cells that were either non activated or activated with 5 μg/ml of HEL (Sigma-Aldrich, 10837059001) or 10 μg/ml AF-488/−647 labelled F(ab’)_2_ Fragments of donkey anti-mouse IgM (Jackson ImmunoResearch 715-546-020/715-606-020), were placed in 300 μl of imaging buffer (10% FCS in PBS) on coverslip wells and left in the incubator (5% CO_2_, +37°C) for 15 minutes. The cells were fixed with 50:50 methanol-acetone at −20°C for 20 min, permeabilised with acetone for 5 minutes at −20°C, and blocked (5% donkey serum in PBS) for 1-2 h at RT. The cells were stained with primary antibodies in PBS supplemented with 5% bovine serum albumin overnight in +4°C and proceeded as above.

For immunostainings, anti-alpha-Tubulin AF488/−647 (DM1A, Merc Millipore, Sigma-Aldrich 16-232/05-829-AF647) was used at the dilution of 1:150 and anti-Golga3 (Sigma-Aldrich HPA040044) was used at 1:150. Anti-HEL antibody (F10, mouse IgG1) was a kind gift from Prof Facundo Batista (the Ragon Institute of MGH, MIT and Harvard, USA). Secondary antibodies antimouse IgG1 AF488/−647 (Jackson ImmunoResearch 115-545-205/115-605-205) and anti-Rabbit IgG AF555 (Invitrogen A-31572) were used at 1:500. Images were deconvoluted with Huygens Essential version16.10 (Scientific Volume Imaging, The Netherlands, http://svi.nl), using automated algorithms and thresholded optimally, yet consistently. Particle analysis was done with Huygens Essential. In Golga3-particle analysis, intensity values of < 5% of the maximum were considered as background. In Golga3-particle analysis, the tubulin channel was used to define cell outlines in 3D that were then shrank by 0.27 μm, corresponding to 1 pixel along Z-axis. Intensities inside the 3D selection were cleared, resulting in the remaining hollow sphere comprising the cell periphery only. Peripheral Golga3 particles were then automatically analyzed. Colocalization between Golga3 and antigen was analyzed with Huygens Essential, also from the peripheral spheres, using optimised, automatic thresholding.

### Statistics and illustrations

Graphs and statistics were prepared with GraphPad Prism (GraphPad Software, La Jolla, CA). Statistical significances were calculated using unpaired or paired Student’s t-test assuming normal distribution of the data, or repeated measures 2-way ANOVA. Statistical values are denoted as: *P<0.05, **P<0.01, ***P<0.001, ****P<0.0001. Illustrations were created with BioRender. Figure formatting was undertaken in Inkscape v.092.2.

## Author Contributions

L.O.A., A.V.S., S.H-P., M.R., V.Š., and P.K.M. conceived and designed the analysis. L.O.A., A.V.S., S.H-P., M.R., B.T-G., D.M.C., M.Ö.B., and V.Š. collected the data. L.O.A., A.V.S., S.H-P., M.R., B.T-G., D.M.C., V.Š., A.V.S., P.P and P.K.M. contributed or performed the data analysis. L.O.A., A.V.S., S.H-P., M.R., B.T-G., and P.P. visualised the data. L.O.A., A.V.S., S.H-P., M.R., B.T-G., and P.K.M. wrote the paper.

## Acknowledgments

We thank Laura Grönfors, Johanna Rajala and Citarra Burrows for technical help. Mass spectrometry analysis was performed at the Turku Proteomics Facility, University of Turku and Åbo Akademi University. Anne Rokka and the staff of the proteomics facility is acknowledged for their help in mass spectrometry analysis. Microscopy and flow cytometry were performed at Turku Bioscience Cell Imaging and Cytometry (CIC), supported by Turku Bioimaging and Euro-Bioimaging. Their personnel is thanked for their generous help and expertise. Biocenter Finland and the InFLAMES Flagship Programme of the Academy of Finland are acknowledged for providing research infrastructures. This work was supported by the Academy of Finland (grant ID: 25700, 296684 and 307313 to P.K.M.; 286712 to V.Š.; 337530 to InFLAMES), Sigrid Juselius and Jane and Aatos Erkko foundations (to P.K.M.), Finnish Cultural foundations (to S.H-P. and V.Š.), Turku Doctoral Programme in Molecular Medicine (TuDMM) (to L.O.A., M.R. and S.H-P.), Magnus Ehrnrooth Foundation (to P.K.M. and A.V.S.) and Turku University foundation (to L.O.A., M.R. and S.H-P.).

## Supplementary Information

- Supplementary Figures 1–5
- Supplementary Files 1-3

**Supplementary File 1. Mass spectrometry data from raft-APEX2 expressing A20 B cells**. A) Proteins identified with high confidence in the whole data set. B) List of proteins exclusively identified in resting cells. C) Proteins exclusively identified in cells activated with anti-IgM surrogate antigen.

**Supplementary File 2. List of proteins defined as lipid raft-resident in B cells**.

**Supplementary File 3. Differential expression analysis comparing resting and activated B cells**. A) List of proteins identified that were used for the differential expression analysis. B) List of proteins that show significant enrichment (adjusted p-values ≤ 0.05) at any given conditions and have log2 fold change ≥ 1.5.

**Supplementary File 4. Identified proteins linked to B cell activation and endocytosis**. A) List of proteins identified within the GO term “B cell activation processes” (GO:0042113). B) List of proteins identified that within the GO term “Endocytosis processes” (GO:0006897).

**Figure S1:**
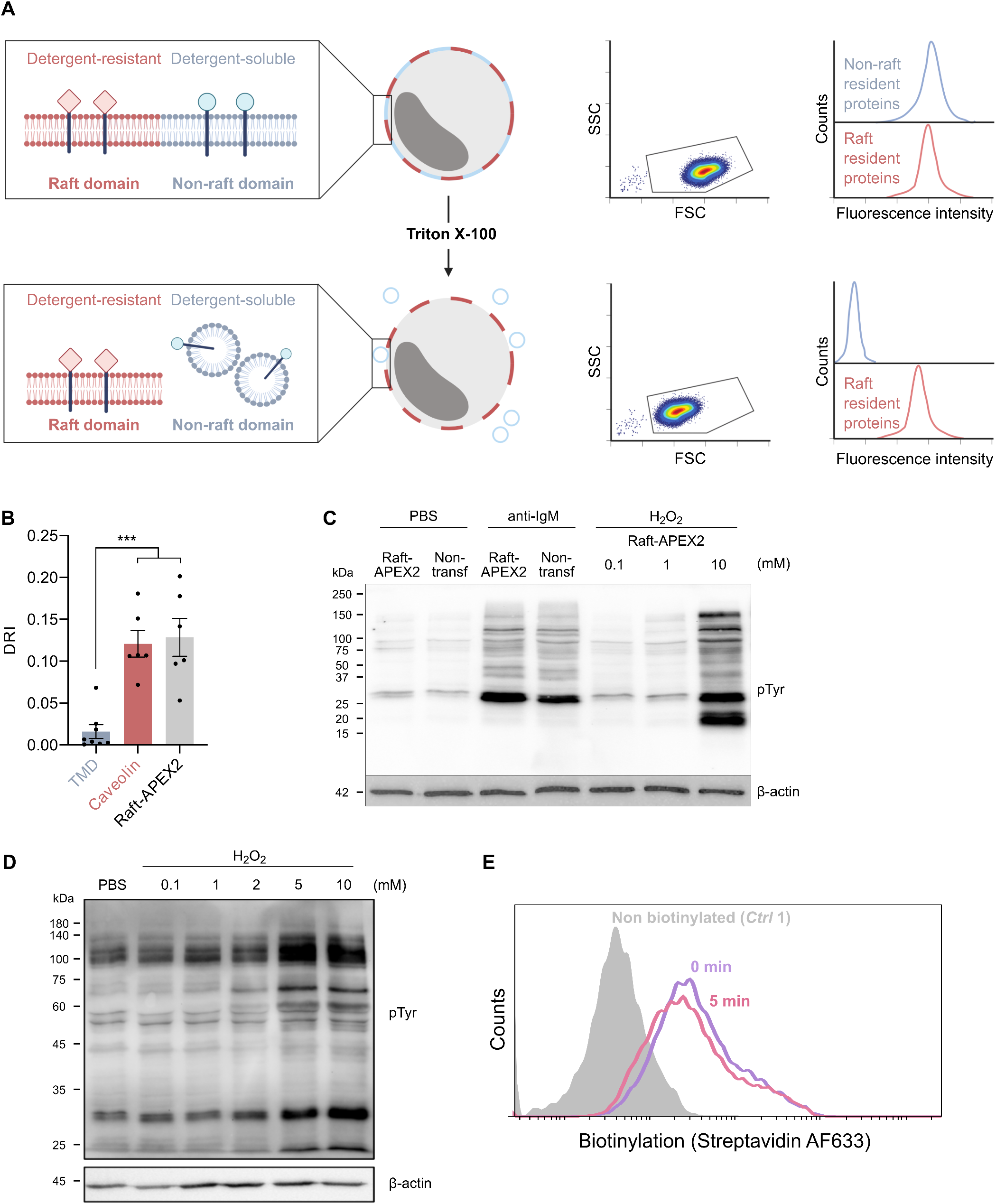
Raft-APEX2 construct locates in the detergent-resistant membrane domains. Related to Fig. 1. A) A schematic representation of flow cytometry assay to detect lipid raft association of membrane proteins by analysis of detergent resistance. Upon treatment with TritonX-100 detergent, the proteins within lipid rafts are retained, whereas non-raft domains are dissolved. B) A20 D1.3 B cells were transfected with raft-APEX2, lipid raft marker (caveolin-1-RFP) or non-raft marker (TMD-GFP). The fluorescence of the markers was measured before, and after subjecting the samples to 0.1% Triton X-100 and the detergent resistance index was calculated as described in Materials and Methods (n = 6-8 independent experiments; unpaired t-test, *** < 0.001). C) The original immunoblot that is shown rearranged in the main Figure 1 D and E. Raft-APEX2 expressing A20 D1.3 B cells and the parental A20 D1.3 (non-transfected) cells were treated with 0, 0.1, 1 and 10 mM H2O2, or 10 μg/ml anti-IgM F(ab’)2 fragments. Cells were lysed and subjected to Western blotting. The membranes were probed with HRP-anti phospho-Tyrosine antibodies and anti-*β*-actin as a loading control. D) An immunoblot like in (C), using Raft-APEX2 expressing A20 D1.3 B cells with 0, 0.1, 1, 2, 5 and 10 mM H2O2 for 1 min. E) Raft-APEX2 A20 D1.3 B cells were supplemented with biotin-phenol, activated (red line) or not with (violet line) F(ab’)2 fragments of anti-IgM antibodies for 5 min, and the biotinylation was triggered or not (grey line) by adding 1 mM H2O2 for 1 min. Cells were fixed with 4% PFA, permeabilised, stained with AF633-labelled streptavidin and analyzed with flow cytometry.

**Figure S2:**
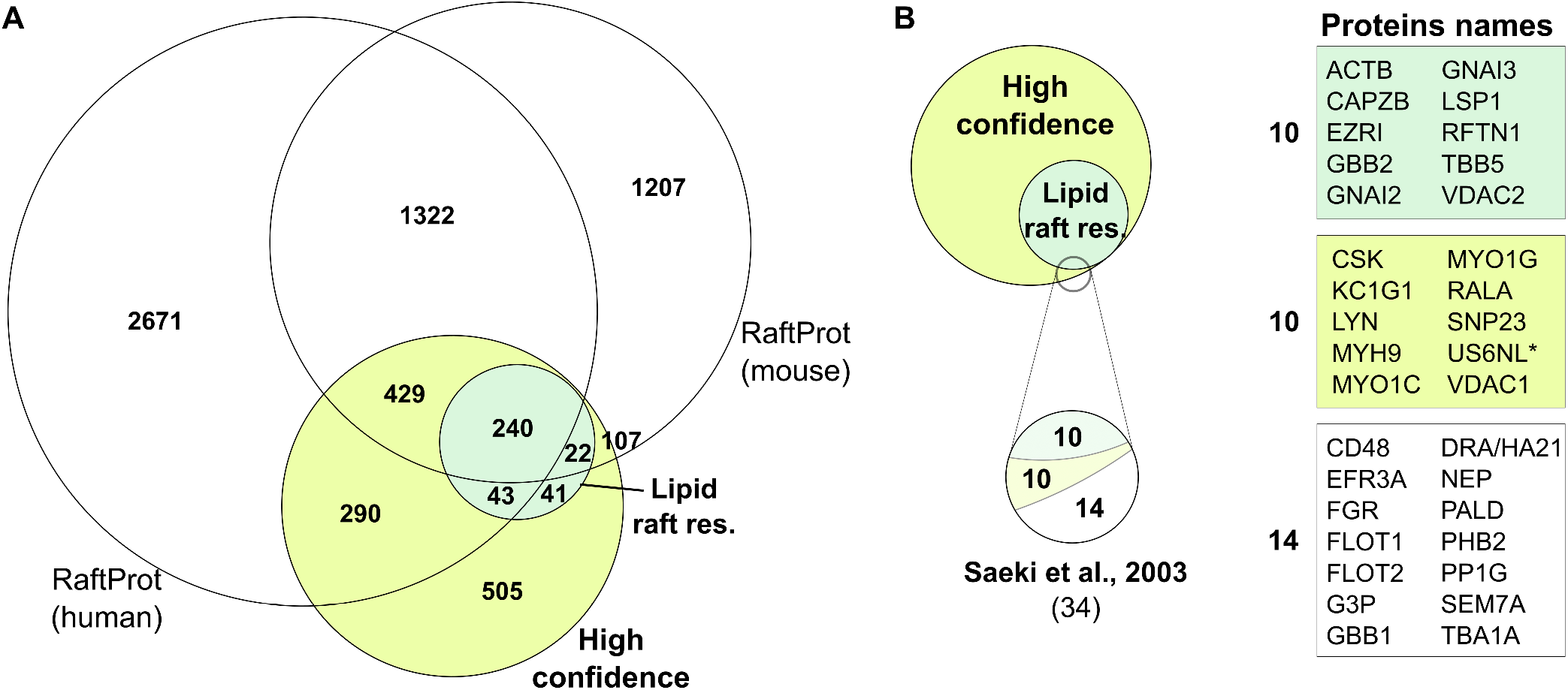
Lipid raft-resident proteins in B cells. Related to main Fig. 3. **A)** A Venn diagram showing intersection between the data obtained in this study to both mouse and human raft proteins in RaftProt database. **B)** A Venn diagram showing intersection between the data obtained in this study to lipid raft proteins identified in Raji B cells by Saeki et al. (*Saeki et al*., 2003)

**Figure S3:**
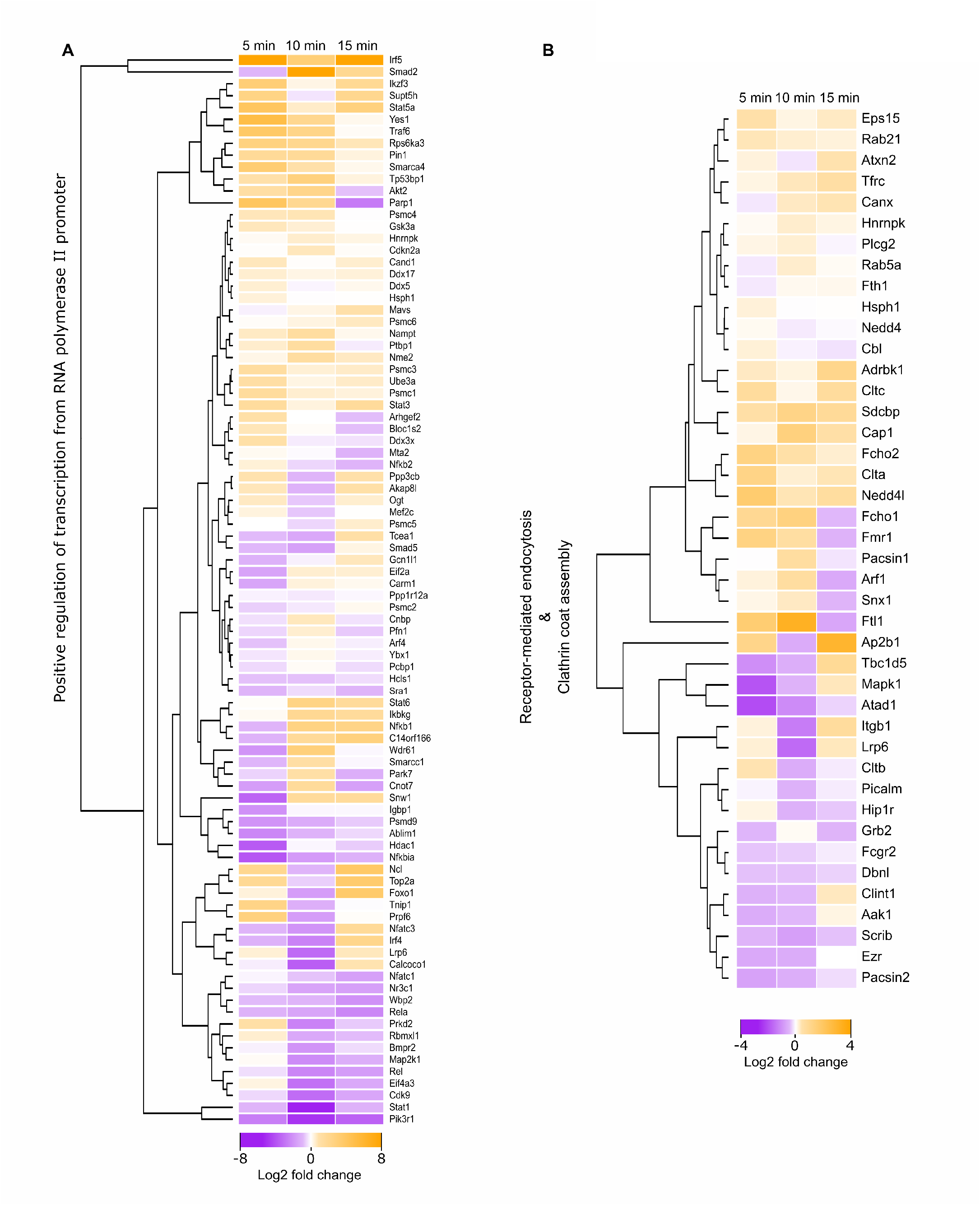
Heatmaps. **A)** Heatmap of the intensity changes of proteins with GO term “positive regulation of transcription from RNA polymerase II promoter”. **B)** Heatmap of the intensity changes of proteins with GO term “receptor-mediated endocytosis” and “clathrin coat assembly”.

**Figure S4:**
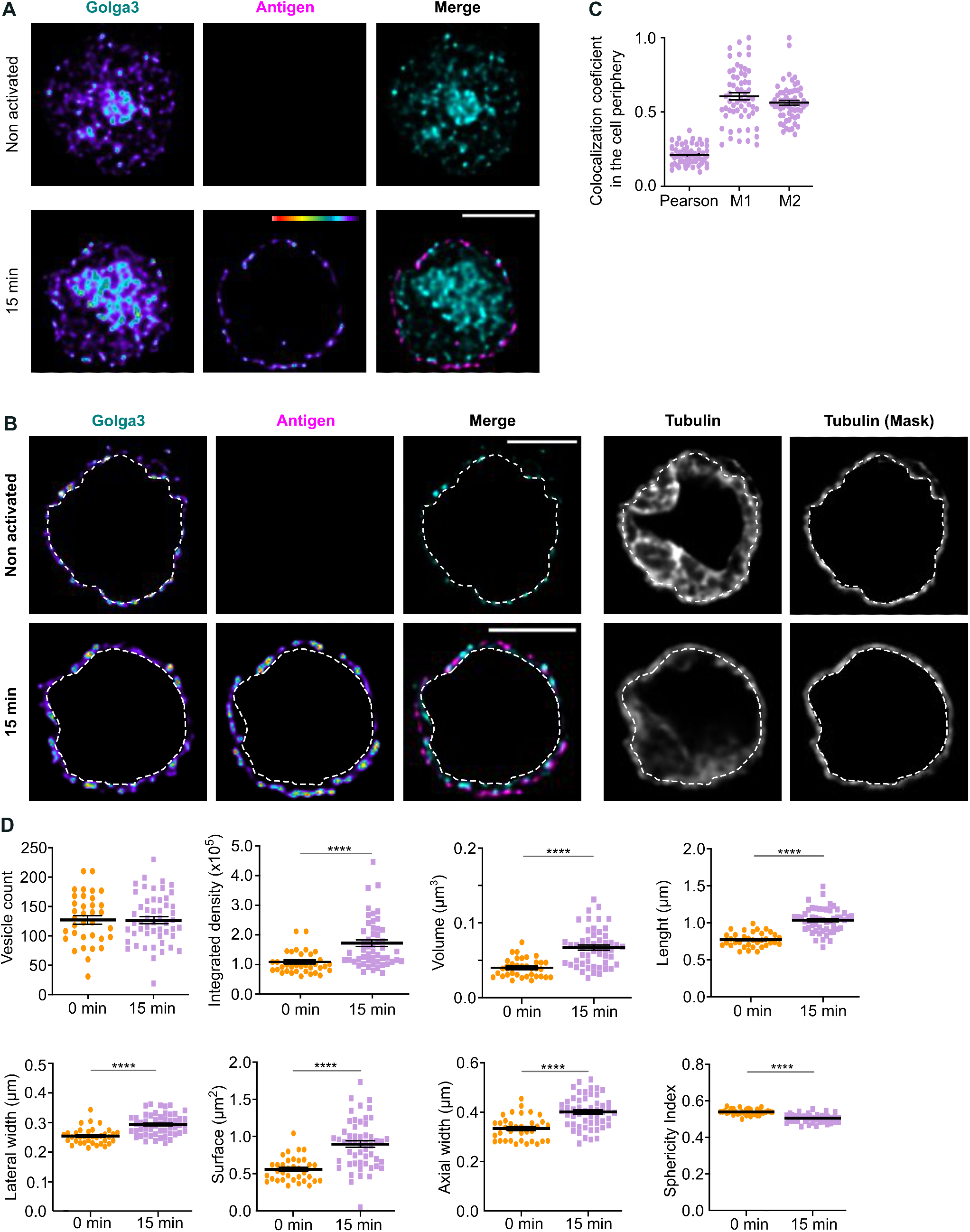
Translocation of Golga3 to the cell periphery upon BCR activation and colocalization with antigen. A) Human Raji D1.3 B cells were let to adhere on CellTak-coated microscopy coverslips, activated (lower panel) or not (upper panel) with 10 μg/ml Alexa Fluor®-labelled F(ab’)2 fragments of anti-IgM antibodies (antigen, pseudocolour in single-channel image and magenta in merge) for 15 min, fixed and permeabilised and subjected to immunofluorescence analysis with anti-Golga3 (cyan) antibodies. Cells were imaged with spinning disc confocal microscopy. B) A 3D mask (dotted line) was generated from the cytosolic tubulin signal, and the inside region of the mask was cleared from signal in all channels. Single confocal planes from deconvoluted images of representative cells (same as in A), showing the filtering, are shown. Scale bar 5 μm. The resulting peripheral Golga3 signal was processed for colocalization analysis with antigen (AF647-anti-IgM F(ab’)2) (C) as well as 3D particle analysis (D) were carried out with Huygens. C-D) The colocalization of antigen and Golga3 (C) as well as the Golga3 vesicles (D) were analyzed from images deconvoluted at the antigen and Golga3 channels, exclusively from the cell periphery, with the filtering illustrated in B. C) The levels of colocalization were measured with Pearson’s correlation coefficiency (0.21) and Manders’ overlap coefficiency (M1: 0.61, M2: 0.56). D) The intensity, volume and length of the peripheral Golga3 vesicles are shown, as well as the vesicle count per cell, lateral and axial width (μm), sphericity index (1=full sphere) and surface area (μm2). Data is shown as mean ± SEM from ≥ 35 cells from three independent experiments. ****: p < 0.0001

**Figure S5:**
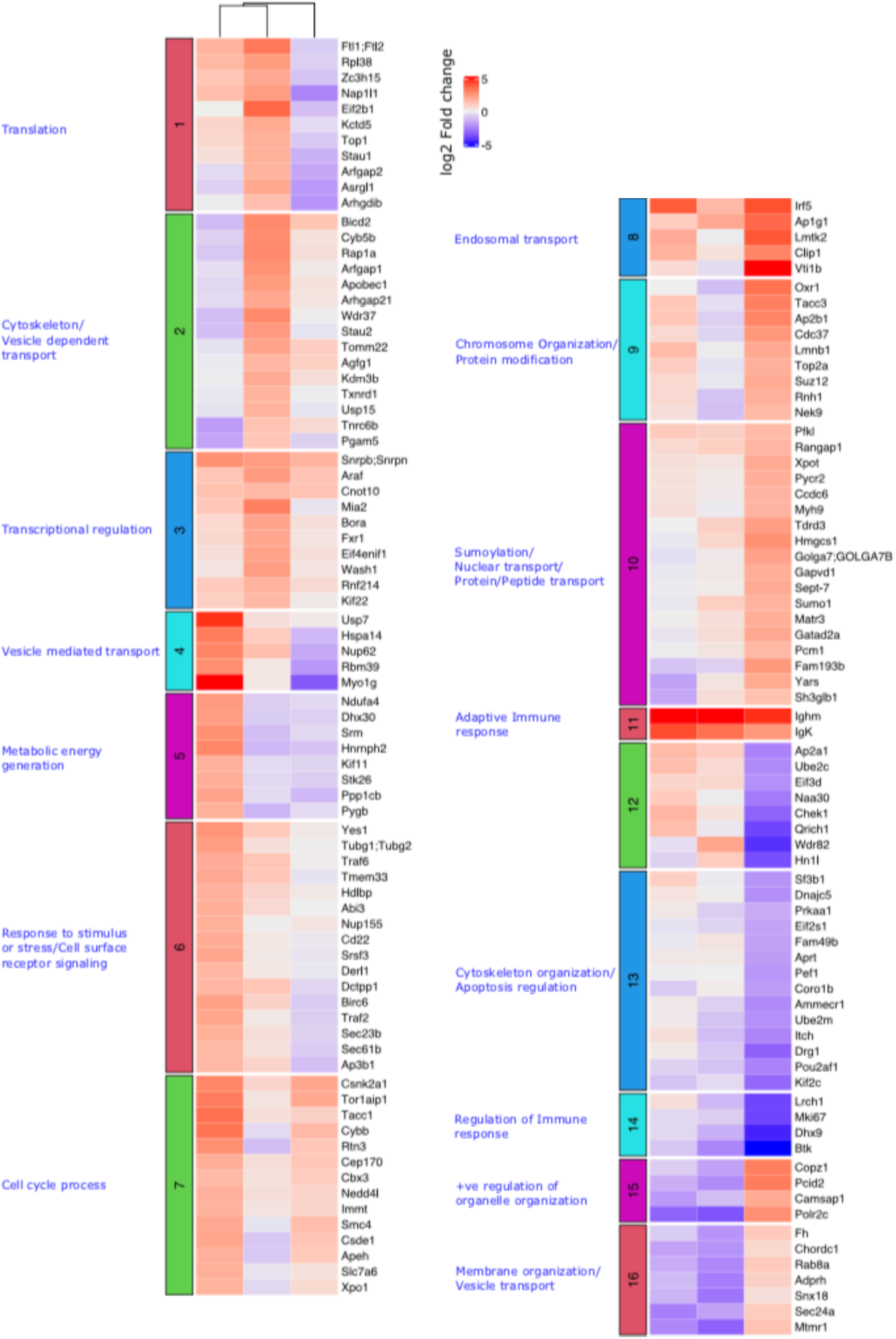
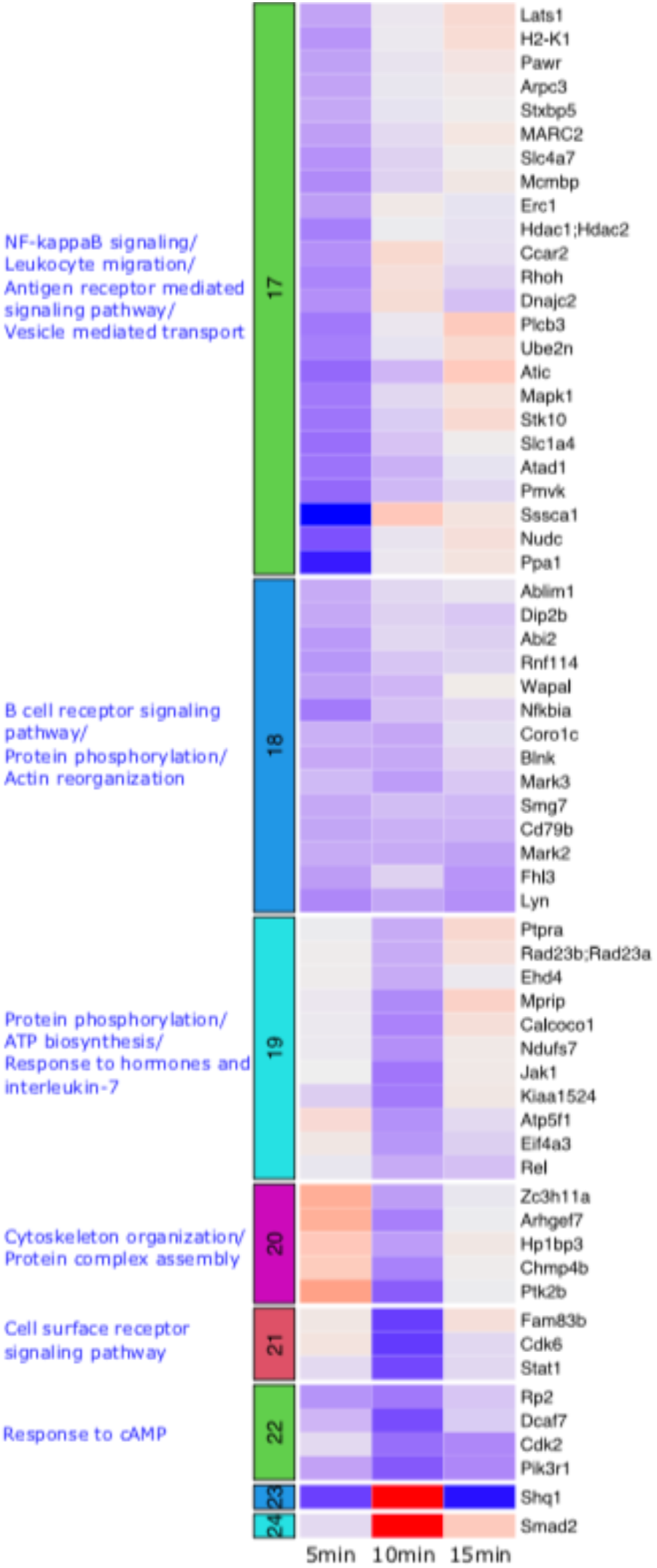
Heat map showing k-means clustering of fold-changes of combined differentially expressed proteins in any of the time points. The purple bands indicate low expression levels while the red bands indicate high expression levels. The optimal number of clusters, k, was determined to be 24 prior to performing k-means clustering. The associated gene ontology (GO), biological processes (BP) of the proteins in each cluster were then determined using DAVID.

**Figure S6:**
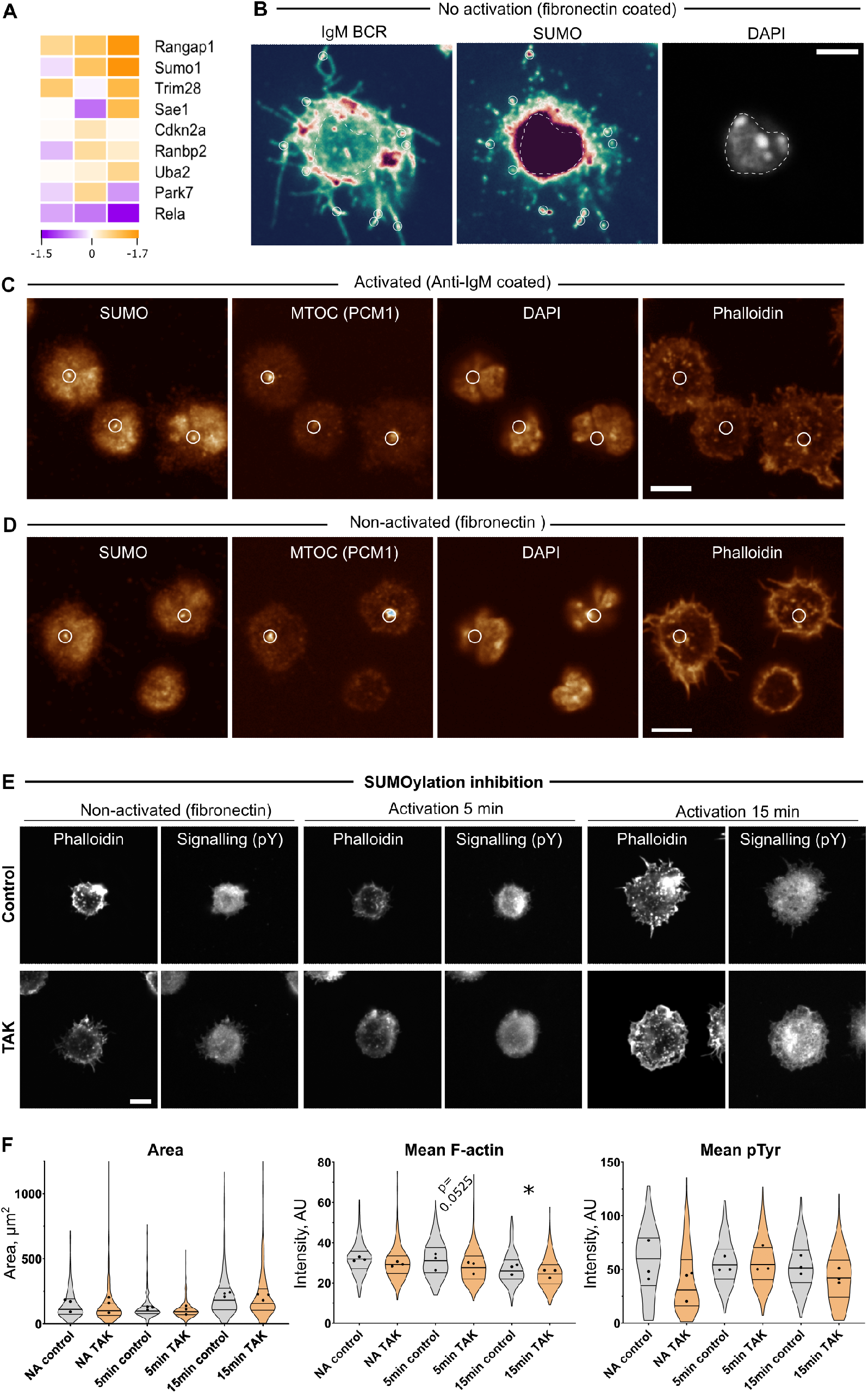
A) Heatmap showing the expression level of proteins associated with SUMOylation in our dataset based on gene ontology (biological process). The purple bands indicate low expression levels while the yellow/orange bands indicate high expression levels. B) SDCM images of non-activated A20 D1.3 cells illustrating the colocalization of the BCR and SUMO (indicated by white circles in the two first panels) and the nucleus (dotted outline in the three panels). Cells were plated on fibronectin, fixed and stained using Donkey anti-mouse IgM F(ab’)2, anti-Sumo1 antibody and DAPI. C-D) Representative images of activated (C) and non-activated (D) A20 D1.3 cells showing the colocalization of SUMO and MTOC (indicated by the white circle). Cells were plated on fibronectin or anti-IgM F(ab’)2 coated surface, fixed and stained with anti-Sumo1 antibody, Anti-PCM1 antibody, DAPI and phalloidin for F-actin. E-F) A20 D1.3 cells were treated with 25 *μ*M TAK-981, to inhibit SUMOylation, or DMSO as control. Cells were plated on fibronectin for non-activatory conditions or on anti-IgM F(ab’)2 for activation for 5 or 15 min, fixed and stained with anti-Phosphotyrosine antibody and phalloidin for F-actin (E). F) Quantification of TAK-981 inhibited and control cells in D. Area of the cell spreading was analyzed by thresholding the phalloidin channel at the cell-surface interface. F-actin and pTyr mean intensity values were measured in the thresholded area. The violin plots represent the distribution of the population (all analyzed cells, >100 cells per experiment and per condition) and the dots represent the means of individual experiments (n=3). All the samples were imaged with spinning disc confocal microscopy, individual slices are shown. Data is shown as violin plots and mean ± SEM of all cells overlaid with individual mean values of three independent experiments (70-100 cells per experiment). Paired t-test on mean values of each experiment (n = 3) was performed. *: p < 0.05. Scale bars = 5 *μ*m.

